# A CDKB/KRP/FB3 cell cycle core complex functions in rice gametes and zygotes

**DOI:** 10.1101/2022.07.12.499798

**Authors:** Hengping Xu, Laura Bartley, Marc Libault, Venkatesan Sundaresan, Hong Fu, Scott Russell

**Affiliations:** Department of Microbiology and Plant Biology, University of Oklahoma, Norman, OK 73019, USA; Institute of Biological Chemistry, Washington State University, Pullman, WA 99164, USA; Department of Agronomy and Horticulture, University of Nebraska-Lincoln, Lincoln, NE 68503, USA; Department of Plant Biology, University of California, Davis, CA 95616, USA

**Keywords:** Cell cycle, coordinate inhibition, core complex, F-Box protein mediated degradation, Kip-Related Protein, *Oryza sativa*, rice zygote, seed formation

## Abstract

The cell cycle controls division and proliferation of all eukaryotic cells and is tightly regulated at multiple checkpoints by complexes of core cell cycle proteins. Due to the difficulty in accessing female gametes and zygotes of flowering plants, little is known about the molecular mechanisms underlying initiation embryogenesis despite the crucial importance of this process for seed crops. In this study, we reveal four levels of factors involved in rice zygotic cell cycle control and characterize their functions and regulation. Protein-protein interaction studies, including within zygote cells, and *in vitro* biochemical analyses delineate a model of the zygotic cell cycle core complex for rice. In this model, CDKB1, a major regulator of plant mitosis, is a cyclin (CYCD5)-dependent kinase; its activity is coordinately inhibited by two cell cycle inhibitors, KRP4 and KRP5; and both KRPs are regulated via F-box protein 3 (FB3)-mediated proteolysis. Supporting their critical role in controlling the rice zygotic cell cycle, mutations in KRP4, KRP5, and FB3 result in the compromised function of sperm cells and abnormal organization of female germ units, embryo and endosperm, thus significantly reducing seed-set rates. This work helps reveals regulatory mechanisms controlling the zygotic cell cycle toward seed formation in angiosperms.

## Background

Rice provides nearly half of daily human nutrition, and rice’s outstanding genetic and genomic resources render it a model monocotyledonous plant. Therefore, understanding the mechanisms underlying the initiation of rice embryogenesis may improve cereal production for the increasing global population (Godfray et al. 2010).

Seeds of monocot plants such as rice consist of embryo and endosperm, derived from double fertilization. The embryo is a diploid embryonic structure with genetic content derived roughly evenly from the sperm and egg; the endosperm is a triploid fusion product of a sperm (n) and a maternal central cell (2n), providing nutrition for the embryo during seed development and germination (Russell 1992; Hu 1998; Dresselhaus et al. 2016; Sprunck 2020). The zygote is the founder cell of the embryo. Its first division forms a smaller cytoplasm-rich apical cell and a larger vacuolated basal cell, which then divides at multiple planes and develops into the pro-embryo and ultimately the embryo (Khanday and Sundaresan 2021).

Embryogenesis is the product of coordinated events of cell division that result in the formation of a mature embryo. The cell cycle needs to be re-established in every generation of sexually reproducing organisms within the nascent zygote. The progression through the cell division phases [gap 1 (G1), DNA synthesis (S), gap 2 (G2), and mitosis (M)] is tightly controlled by multiple regulatory proteins at each checkpoint, especially in the transitions from G1 to S and G2 to M. Cell cycle control coordination failures can be lethal, leading either to embryo abortion or to development of tumors (Williams and Stoeber 2012; Otto and Sicinski 2017).

Across eukaryotes, Cyclins (CYC) and Cyclin-Dependent Kinases (CDKs) constitute the central core cell cycle control proteins. Flowering plants possess a large set of CDKs and CYCs that allowing numerous core complex combinations (Stals and Inze 2001; De Veylder 2007; Van Leene 2011; Cheng et al. 2013; Dante 2014; Pedroza-Garcia 2016; Wang 2004; La 2006; Dudits 2007; Hu et al. 2010; Ma et al. 2013; Lin et al. 2014). Furthermore, both CDK and CYC are conjugated with a third level partner, an Inhibitor of Cyclin-Dependent Kinase (ICKs), which inhibits CDK activity (Dudits 2016; Lin et al. 2014), and in plants are called Kip-Related Proteins (KRPs; Kip, a mammalian tumor suppressor). KRPs are nuclear proteins of 100 to 300 amino acid residues, with 2 motifs at the C-terminal that interact with CDK and CYC (Stals and Inze 2001; Wang 1997; De Veylder et al. 2001). In Arabidopsis, plant sperm cell formation is controlled by two cell cycle inhibitors, AtKRP6 and AtKRP7, which are degraded by the FB Like 17-associated SCF complex (Kim et al. 2008). Rice has six KRPs, most of which play a critical role in rice seed development and vegetative growth. For instance, KRP1 functions in rice plant growth and seed formation (Barrôco et al. 2006). Overexpression of KRP1 and KRP2 results in smaller seeds (Ajadi et al. 2020). KRP3 controls rice syncytial endosperm formation (Mizutani et al. 2010), and over-expressing KRP4 results in abnormal rice growth and reduced seed-set rate (Yang et al. 2011). But, to date, no KRPs have been specifically connected to rice zygotic cell cycle control.

To enter the S or M-phase, the inhibitory KRPs must be removed so that CDKs can be activated. Members of the F-box protein (FBP) family play a central role by mediating the degradation of KRPs. Numbers of FBPs vary greatly among species (Kipreos and Pagano 2000; Xu et al. 2009; Zhang S et al. 2019; Zhang X et al. 2019). In rice, for example, 678 FBPs have been predicted. Across eukaryotes, FBPs typically associate with two other factors to form an SCF complex (Skp1-Cullin1-F-box) (Kipreos and Pagano 2000; Pagano et al. 1995; Bai et al. 1996; Schulman et al. 2000; Zheng et al. 2002; Malik et al. 2020), which acts as an E3 ligase that targets protein ligands for proteolysis via the 26S proteasome pathway (Zhang et al. 2019).

Due to the inaccessibility of the zygote, which is deeply embedded within multiple layers of tissue, no core complex (CYC-CDK and KRP) has been identified that controls rice zygotic division. Here, we report a rice cell cycle core complex consisting of CYCD5 and CDKB1 that is coordinately arrested by two inhibitory proteins, KRP4 and KRP5, and, in turn, activated by FB3-mediated proteolysis. This complex is likely involved in rice zygotic cell cycle control.

## Results

### Rice KRP4 and KRP5 are preferentially expressed in egg cells and zygotes

Cell cycle control core complexes include three components: CDKs, CYCs and KRPs. Interactions among these components have been comprehensively studied in Arabidopsis (Leene et al. 2011). However, in rice, we have limited knowledge of these interactions. Particularly, little is known of their function in female gametes and zygotes of Arabidopsis or other flowering plants. Due to the well-known difficulty in accessing these single cells that are deeply embedded within multiple layers of tissue, there is no research on the functions of cell cycle core complex in plant female gametes or zygotes for the initiation of embryogenesis following fertilization. To address this void, we mined the relevant literature (Guo et al. 2007) and genomic and transcriptomic (Anderson et al. 2013; Anderson et al. 2017) databases, and identified 9 *CDK*, 6 *KRP*, and 24 *CYC* genes expressed in rice flowers (Additional file 2: Table S1). Among them, 3 *KRPs* (*KRP1, KRP4* and *KRP5*), 6 *CDKs* (*CDKA1;1, CDKA2;1, CDKB1;1, CDKB2;1, CDKC1;*2 and *CDKE1*), and 10 *CYCs* were cloned for further functional study as candidate core complex factors in rice zygotic cell cycle.

Transcriptome data revealed preferential expression of *KRP4* and *KRP5* in rice egg cells and zygotes, compared to other *KRP* genes (Anderson et al. 2013; Anderson et al. 2017) (Fig. 1A). We further confirmed by RT-qPCR that the expression of *KRP4* and *KRP5* is significantly higher in rice pistil compared to other tissures (leaf, stem, root anther, and lemma/palea, Fig. 1B, C).

**Fig. 1.**
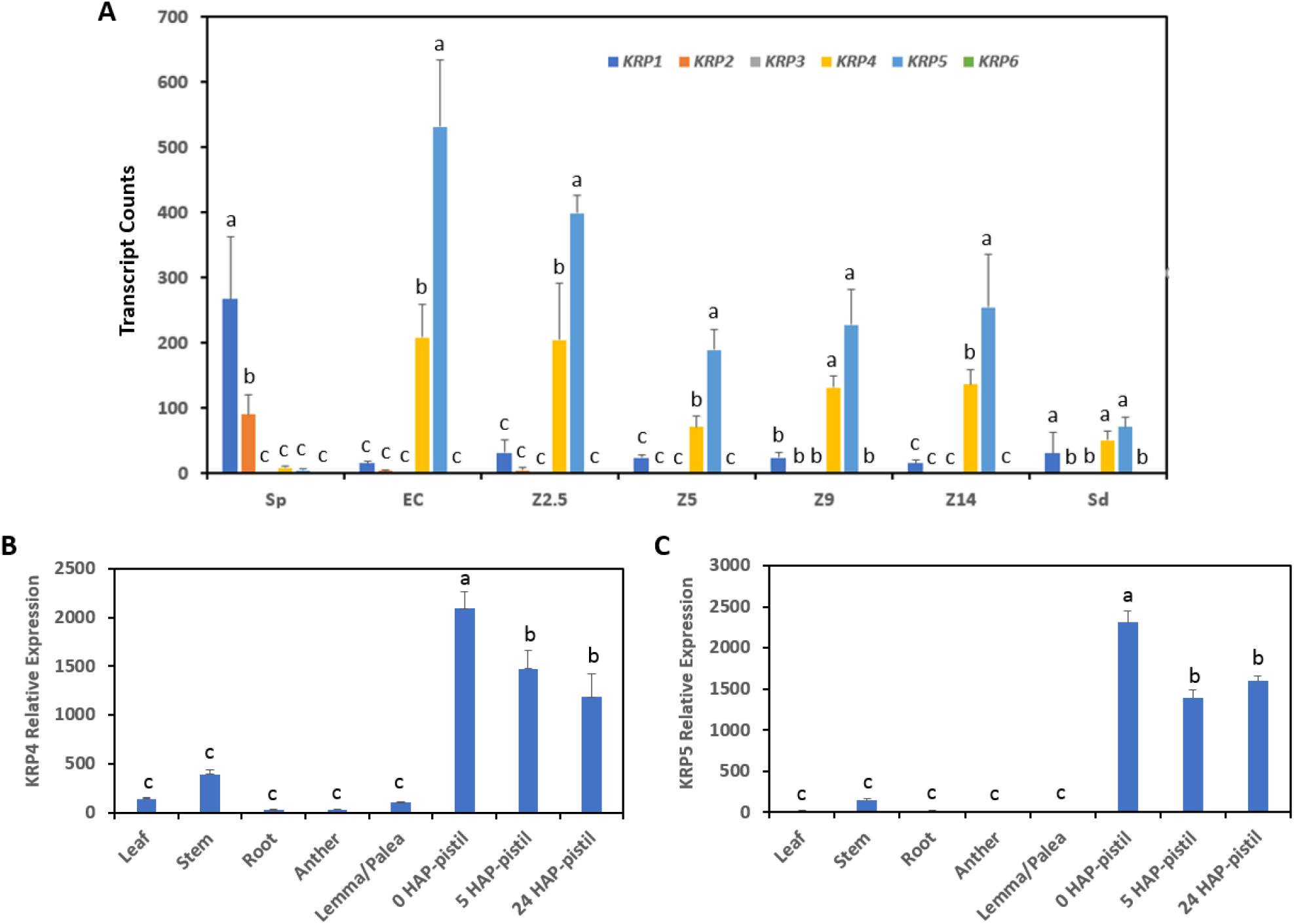
*KRP4* and *KRP5* genes are preferentially expressed in rice egg cells and zygotes. **A** Transcription of 6 rice *KRP* genes in rice egg cells and zygotes, based on RNAseq data (Anderson et al., 2013; Anderson et al., 2017). Sp, sperm cell; EC, egg cell; Z2.5, zygote at 2.5 hours after pollination (HAP); Z5, zygote of 5 HAP; Z9, zygote of 9 HAP; Z14, zygote of 14 HAP; Sd, seedling. **B** RT-qPCR expression profiles of *KRP4* in different organs. **C** RT-qPCR expression profiles of *KRP5* in different organs. Data are means of 3 biological replicates and error bars represent the standard deviation. Significantly different groups are indicted with different letters determined by one-way ANOVA (with Tukey’s post-hoc test, *P* < 0.05).

### Yeast two hybridization (Y2H)-based interactome of cell cycle core complex proteins of rice egg cells and zygotes

To elucidate the proteins potentially composing the cell cycle core complex, we conducted Y2H assays using KRP4 and KRP5 proteins as baits, and CDK and CYC proteins as preys. Using a strict selective media, we detected that KRP5 interacts with all 6 CDKs (CDKA1;1, CDKA2;1, CDKB1;1, CDKB2;1, CDKC1;1 and CDKE1, Fig. 2A). Similarly, KRP4 interacts with all CDKs except CDKE1. Notably, we observed a preferential interaction between KRP4 and CDKC1;2. As expected, truncated KRP4 (KRP4m) and KRP5 (KRP5m), lacking the domain for interaction with CDK, did not interact with any of the six CDKs tested (Fig. 2A).

**Fig. 2.**
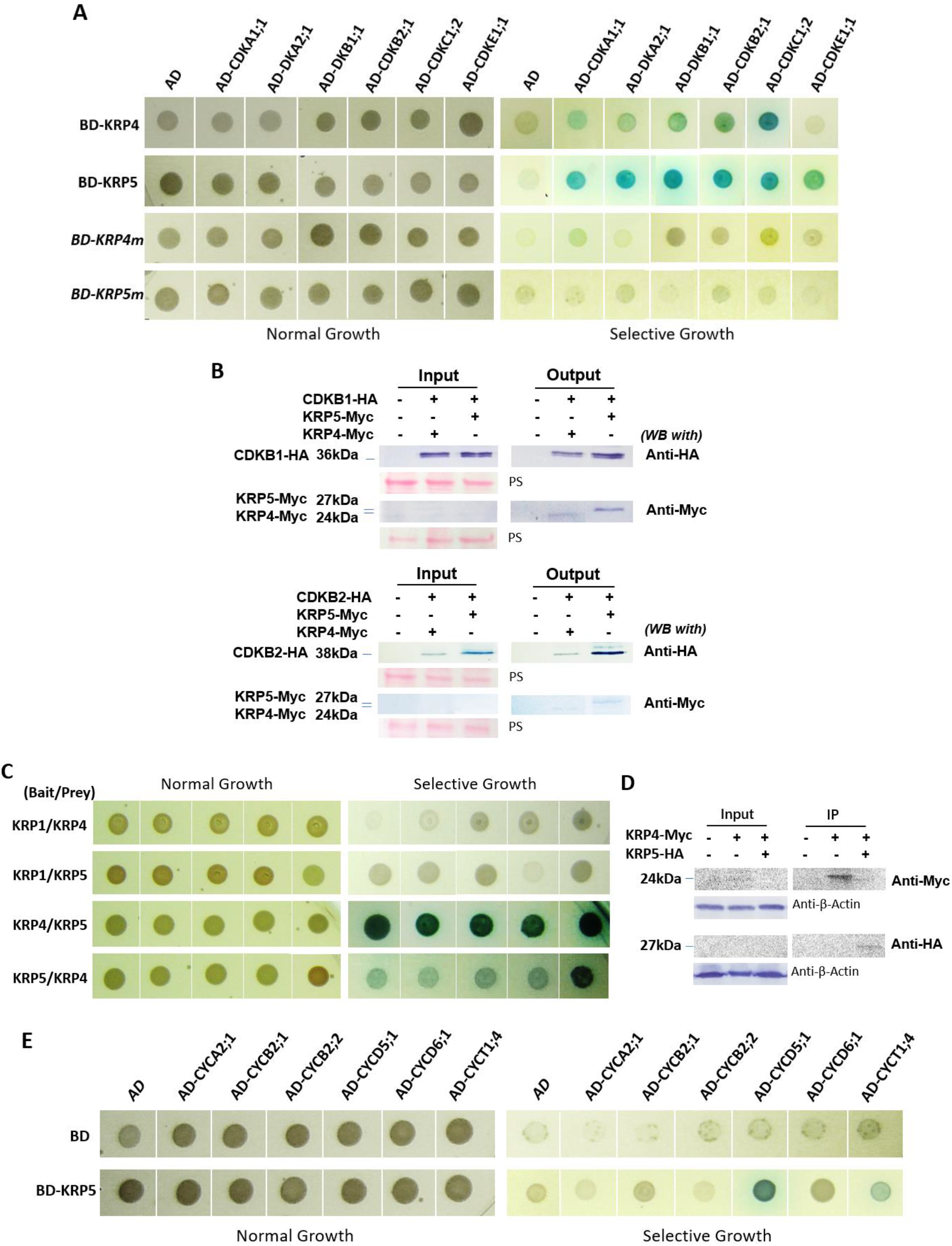
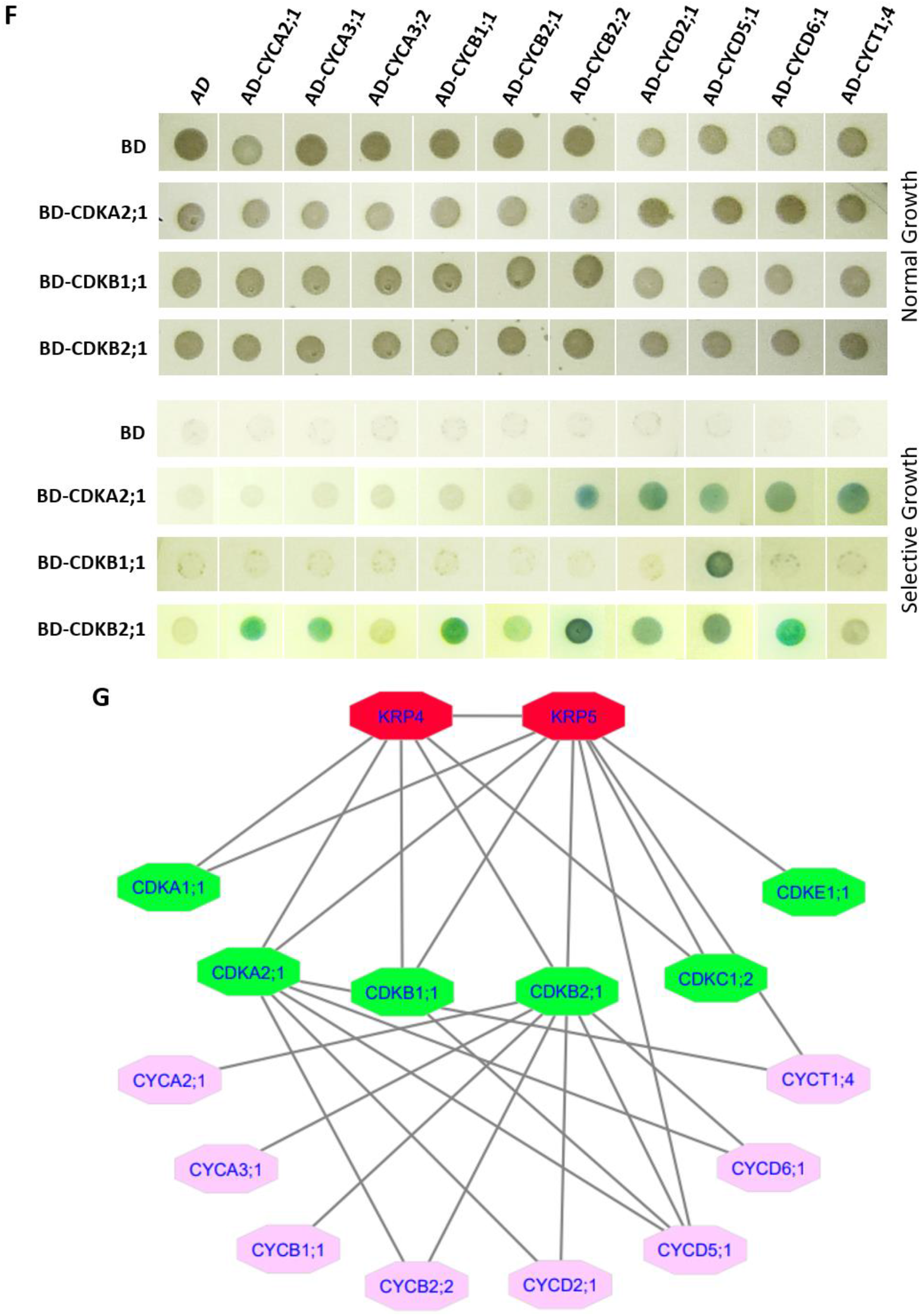
Interactions among rice cell cycle proteins expressed in yeast. **A** Interactions of KRP4 and KRP5 with CDKs. The blue spots on the selective medium indicate interactions of KRP4 or KRP5 with rice CDKs. The prey vector pGADT7 (AD), and both truncated KRP4 and KRP5 (i.e., C-terminal truncations KRP4m and KRP5m) as baits were used as negative controls. Similar results were observed in three independent assays. In A, C, E and F, the normal growth is on DDO (SD-Trp-Leu) and the selective growth is on QDO (SD-Trp-Leu-His-Ade) with X-α-gal and Aureobasidin A (AbA), at 30°C for 3 days. **B** Co-immunoprecipitation (Co-IP) for interactions of KRP4 and KRP5 with CDKB1 or CDKB2 was conducted with Anti-c-Myc antibody using nuclear proteins extracted from the yeast cells harboring CDKB1-HA with KRP4-c-Myc or KRP5-c-Myc (the upper panel) or harboring CDKB2-HA with KRP4-c-Myc or KRP5-c-Myc (the lower panel). Proteins on Western blots were detected with Anti-HA and anti-c-Myc antibodies. Equal loading of protein samples is indicated by Ponceau staining (PS). **C** Y2H was used to detect the interaction (indicated by blue spots) of KRP4 and KRP5. KRP1 was used as the bait for the control. Similar results were obtained from two independent experiments. **D** Co-IP (lower panel) was conducted with Anti-c-Myc for the interaction between KRP4 and KRP5; the Western blots were probed with Anti-HA and Anti-c-Myc and visualized in a chemiluminescent system. Actin detected with Anti-ß-Actin was used as a loading control. **E** Interactions of KRP5 with different rice CYCs (blue spots). KRP5 interacts with CYCD5 and CYCT1:4. A similar outcome was obtained in three separate assays. **F** Interactions of CDKA or CDKB with different rice CYCs (blue spots). CDKB1 interacted only with CYCD5. Three independent assays were performed with similar observations. In E and F, both bait vector (BD) and prey vector (AD) were used as negative controls. **G** Scheme representing Y2H-based interactome of cell cycle core complex components in rice egg cells and zygotes displayed with Cytoscape 3.8.2.

As CDKB proteins are unique to plants and essential for plant cell division during mitosis (Leene et al. 2011; Boudolf et al. 2004; Andersen et al. 2008; Atkins and Cross 2018), we verified the interaction of CDKB1 and CDKB2 with KRP4 and KRP5 in a co-immunoprecipitation assay. Both c-Myc-tagged KRP4 and KRP5 and HA-tagged CDKB1 were precipitated from total yeast nuclear proteins with an anti-Myc antibody [Fig. 2B (upper panel)], demonstrating the interactions of CDKB1 with both KRP4 and KRP5 in yeast nuclei. Similarly, CDKB2 also interacts with both KRP4 and KRP5 [Fig. 2B (lower panel)].

Since both KRP4 and KRP5 interact with most selected CDKs (Fig. 2A & B), we traded their orientation (as bait or prey) in Y2H to assess specificity and also used these constructs to test interactions between the KRPs. Results were similar regardless of orientation, and the KRP proteins strongly interacted with each other (Fig. 2C & D). To our knowledge, this is the first interaction detected between the two putative inhibitory KRP proteins in eukaryotes. Importantly, subsequent experiments verified the coordination between KRP4 and KRP5.

In addition, KRP1, KRP4 and KRP5 were used as baits for interactions with different CYCs. As shown in Fig. 2E, we detected interactions only between KRP5 and CYCD5;1 or CYCT1;4 (weakly). All six rice CDKs were also used as baits to detect potential interactions with ten different rice CYCs. We found that CDKA2;1 interacts with five CYCs (CYCB2;2, CYCD2;1, CYCD5;1, CYCD6;1 and CYCT1;4), CDKB1;1 interacts only with CYCD5;1; and that CDKB2;1 interacts with seven Cyclins (CYCA2;1, CYCA3;1, CYCB1;1, CYCB2;2, CYCD2;1, CYCD5;1 and CYCD6;1) (Fig. 2F). No interaction was detected from the other three CDKs and ten CYCs. All these detected interactions are summarized as Fig. 2G.

### KRP4 and KRP5 coordinately interact with CDKBs to inhibit yeast growth and kinase activity

Figure 3A and B, show that transfection with a single rice cell cycle control protein, either CDKB1, CYCD5, KRP4 or KRP5, does not significantly affect yeast growth. However, co-transformation of CDKB1 (Fig. 3A), or CDKB1 and CYCD5 (Fig. 3B; Additional file 1: Fig. S1), with either KRP4 and/or KRP5, singly or together, significantly reduces the yeast growth rate. This observation suggests that the complexation of CDKB1 (and CYCD5) with KRP4 or KRP5 can negatively regulate the yeast cell cycle. The lower growth rate when both KRP4 and KRP5 are present suggests a cooperative relationship between KRP4 and KRP5 in their interactions with CDKB1 (and CYCD5).

**Fig. 3.**
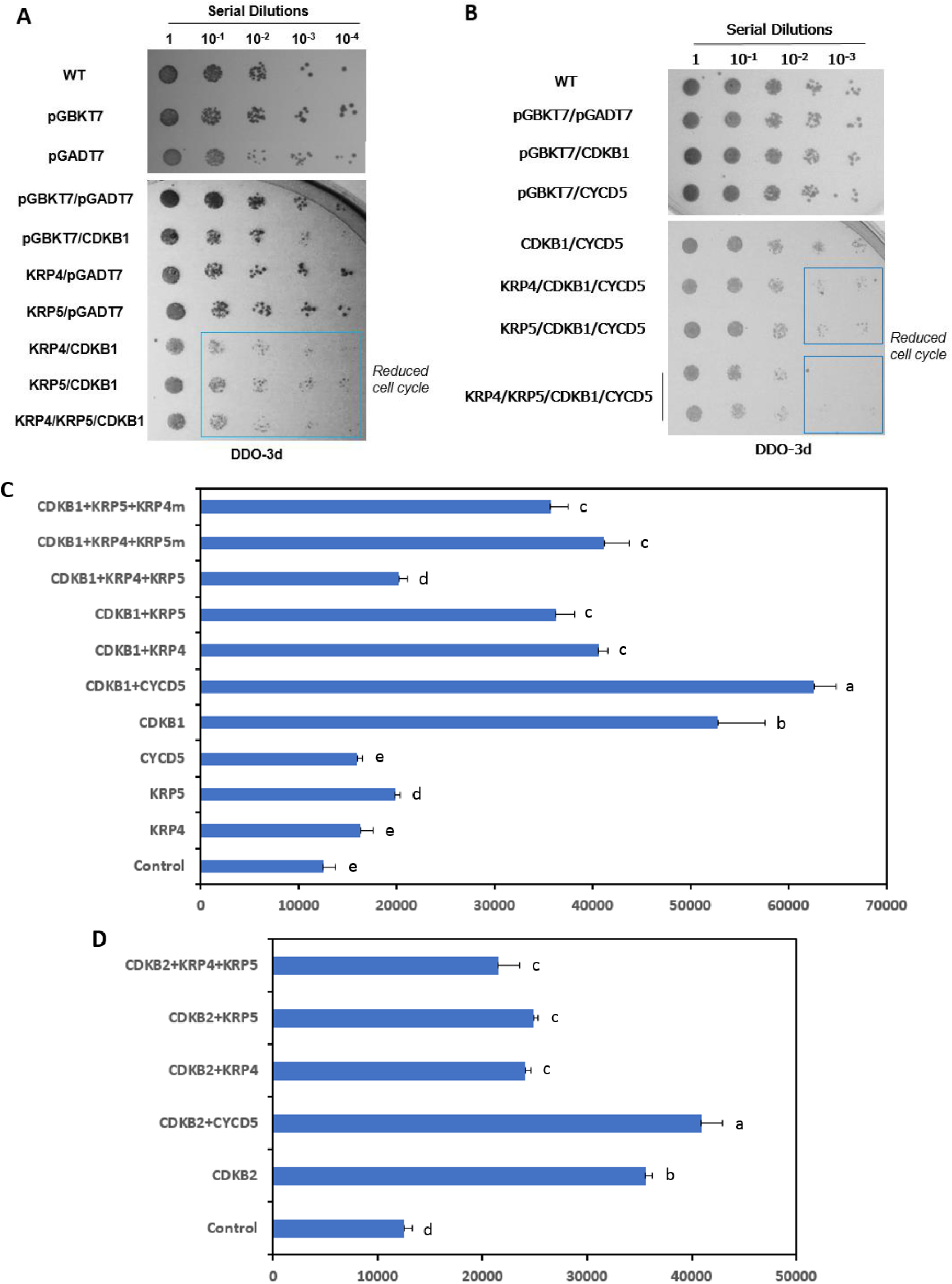
KRP4 and KRP5 coordinately interact with CDKB. **A** Growth of serial dilutions of yeast cells transfected with rice core cell cycle genes indicating the interactions between rice CDKB1 with KRP4 and KRP5. **B** Growth of serial dilutions of yeast cells transfected with core rice cell cycle genes indicating interactions of rice CDKB1-CYCD5 with KRP4 and KRP5. In A and B, BD represents the bait vector, pGBKT7; AD, the prey vector pGADT7. DDO-3d, growth on the medium SD-Trp-Leu for 3 days. Blue boxes demark the inhibition of yeast growth due to the presence of cell cycle inhibitory complexes from rice. Similar results were observed in 3 independent experiments. **C** ADP Sensor™ kinase activity assay showing coordinated inhibitory effect of KRP4 and KRP5 on Kinase Activity of CDKB1. **D** Kinase activity assay showing coordinated inhibitory effect of KRP4 and KRP5 on kinase activity of CDKB2. In C and D, Kinase activity was measured in relative fluorescent units (RFU). – Control: reaction containing no recombinant protein eluted in Co-IP using yeast cells harboring only the empty pGBKT7 and pGADT7 vectors. CYCD5, rice Cyclin D5;1. The activities of CDKB1 (C) and CDKB2 (D) are increased by CYCD5 but maximally decreased by both KRP4 and KRP5. KRP4m, the C-terminally truncated protein encoded of KRP4 (N-terminal -105 amino acids); KRP5m, the C-terminally truncated protein of KRP5 (N-terminal -177 amino acids). Values in C and D are mean ± standard deviation, N=3 biological replicates. Letters indicate significant differences among group means determined by one-way ANOVA analysis (with Tukey’s post-hoc test, *P* < 0.05).

Both rice KRP4 and KRP5 are predicted to function as CDK inhibitors based on sequence homology. We verified their inhibitory function on CDK kinase activity with rice proteins expressed and immunopurified from yeast nuclei. In the assay that detects ADP formation as a readout of kinase activity, base activity of CDKB1 alone was ∼52,797 relative fluorescence units (RFU), which increased by ∼20% upon the addition of CYCD5 (Fig. 3C). On the other hand, CDKB1 activity decreased by 23% and 31% upon incubation with KRP4 and KRP5, respectively. Strikingly, CDKB1 activity dropped by 62% when both KRP4 and KRP5 were added (CDKB1 + KRP4 + KRP5). This additive inhibition effect on the activity of CDKB1 is reversed back by replacing one of the two KRPs with its truncated form (i.e., KRP4m or KRP5m), lacking the CDK-interaction domain. The inhibitory effect of KRP4 and KRP5 was also observed on the activity of CDKB2 (Fig. 3D). The kinase activity of CDKB2, estimated at 35,390 RFU, increased by 15% upon incubation with CYCD5, and decreased by 32% and 30% by incubating with KRP4 and KRP5, separately. CDKB2 activity decreased by 40% upon incubation with both KRP4 and KRP5 (CDKB2 + KRP4 + KRP5), supporting the additive inhibition of KRP4 and KRP5 on the CDKB activity. The above assays demonstrate the coordinate relationship of KRP4 and KRP5 in their inhibition of CDKB activity.

### Bimolecular Fluorescence Complementation (BiFC) shows interactions between core cell cycle proteins in isolated living rice egg cells and zygotes

As shown in Fig. 4A, fluorescent signals (green) that appear with the native promoter-driven BiFC constructs evidence *in vivo* interactions of KRP4 and KRP5 with CDKB1 in rice egg cells and zygotes. These interactions take place within the nucleus as indicated by the overlap of EYFP signal with a mCherry protein fused with a nuclear localization signal (NLS). This result corroborates previous studies showing the nuclear localization of the KRP proteins (Yang et al. 2011). As shown by the positive fluorescent signal in Fig. 4A, both KRP4 and KRP5 interact with CDKB1 in rice egg cells with very similar patterns, consistent with the interactions among CDKB1, KRP4, and KRP5.

**Fig. 4.**
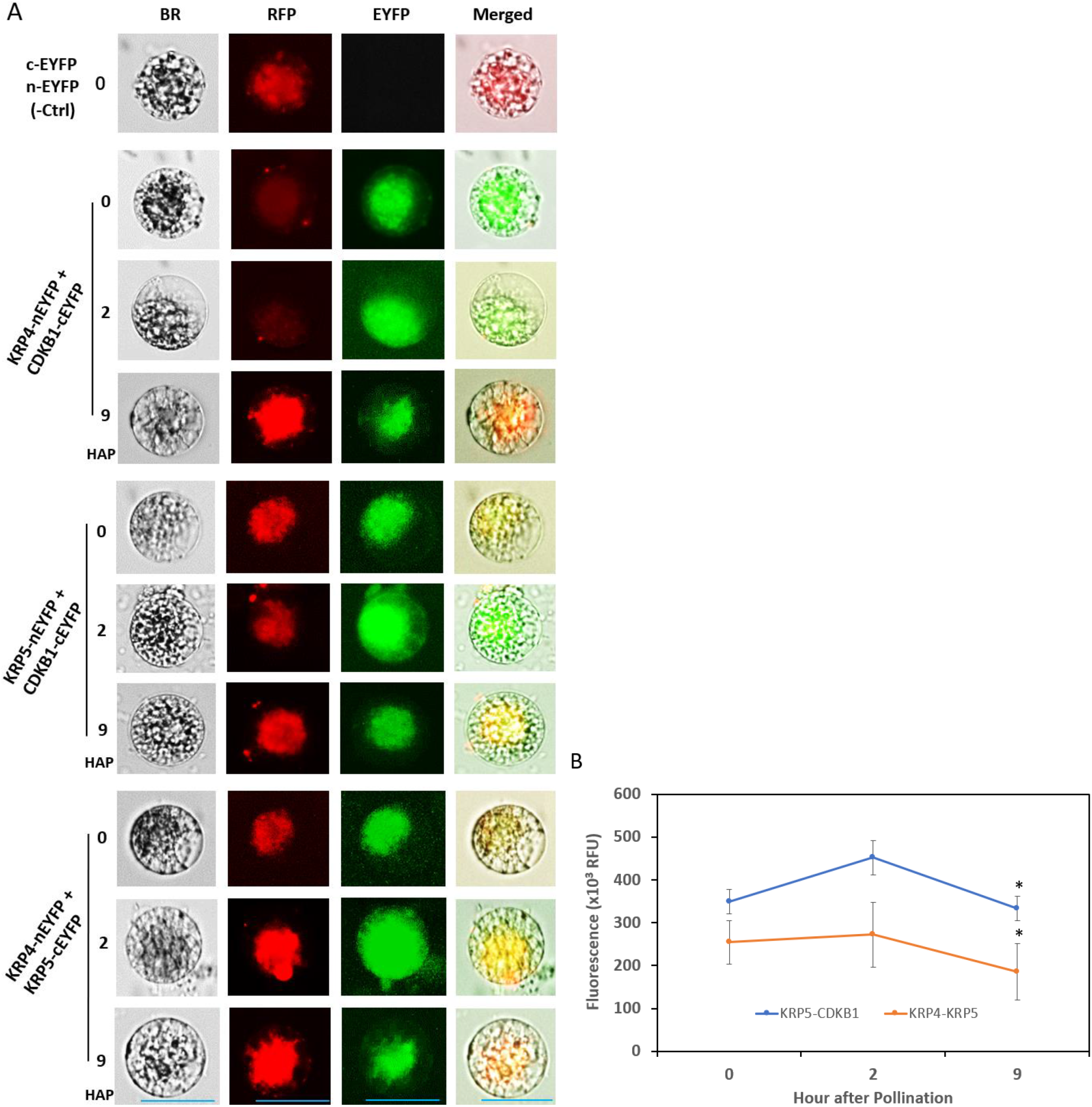
Bimolecular Fluorescent Complementation (BiFC) demonstrates interactions of core cell cycle complex components in rice egg cells and zygotes. **A** Representative images of BiFC showing the interactions of KRP4 with CDKB1 in the egg cell and zygote (harvested at 2 and 9 HAP) transfected with the construct *Ubi*_*pro*_*-KRP4-nEYFP + CDKB1*_*pro*_*-CDKB1-cEYFP, in* 2 biological replicates (N = 3 cells in each); the interactions of KRP5 with CDKB1 in the egg cell (0 HAP) and zygotes (2 and 9 HAP) transfected with the construct *KRP5*_*pro*_*-KRP5-nEYFP + CDKB1*_*pro*_*-CDKB1-cEYFP, in* 3 biological replicates (N = 3-6 cells in each); and the interaction of KRP4 and KRP5 in rice egg cells and zygotes (2 and 9 HAP at harvest) transfected with the construct *Ubi*_*pro*_*-KRP4-nEYFP + KRP5*_*pro*_*-KRP5-cEYFP*. The empty vectors containing *nEYFP* and *cEYFP* were used for the negative control (-Ctrl); BR show cell images under bright field illumination; RFP shows the localization of mCherry_NLS_, where NLS stands for nuclear localization signal; EYFP shows the interaction of the two partner proteins; the merged image is compiled from all three images. In A and B, scale bars: 50 µm. **B** Mean fluorescence from cellular images of BiFC between KRP4 and KRP5 plus those between KRP5 and CDKB1. Each observation had 3 biological replicates (N = 3-6 cells in each). The error bars denote the standard deviation. The * indicates the significant difference of apparent protein interaction levels at 9 HAP vs 2 HAP via one-way ANOVA test (*P* < 0.05).

As in the yeast and kinase activity assays, KRP4 and KRP5 also interact in native promoter-BiFC assays in rice egg cells and zygotes (0, 2 and 9 HAP, Fig. 4B). Remarkably, we noticed that the apparent relative fluorescent units (RFU) of the transfected cells varies with the stages of zygotic development at harvest (Fig. 4B). Compared to the egg cells, the intensity was slightly higher (∼ 6.9%) in the zygotes at 2 HAP, but apparently lower (∼ 32%, *P*<0.05) in zygotes at 9 HAP. Meanwhile, the interaction of KRP5 and CDKB1 with development presents the similar pattern (Fig. 4B). This observation seems fit for a model in which both KRP4 and KRP5 function as CDKB inhibitors as the cell cycle proceeds to the crucial stage: the zygotic division (at ∼12 HAP). However, since the fluorescent intensity observed in BiFC is influenced by different stimuli (Kerppola 2008), the pattern (Fig. 4B) remains to be further confirmed with advanced methods.

### FB3 protein interacts with KRP4 and KRP5

Based on cell cycle regulation in other organisms, we next sought to determine if a rice F-box protein might interact with KRP4 and KRP5, potentially targeting them for degradation. Of the 678 F-box genes annotated in rice, only seven (referred to as *FB1 to FB7* in Additional file 2: Table S3) are expressed in rice gametes and zygotes according to RNAseq data (Anderson et al. 2013; Anderson et al. 2017). Among them, *FB3* is preferentially expressed in rice egg cells and zygotes (Additional file 1: Fig. S2). We confirmed this result by RT-PCR in which we detected the highest transcript abundance in the rice pistil (0 HAP) compared with other organs (Fig. 5A). Moreover, Y2H assays revealed that the FB3 protein interacts moderately with KRP4 and strongly with KRP5 (Fig. 5B).

**Fig. 5.**
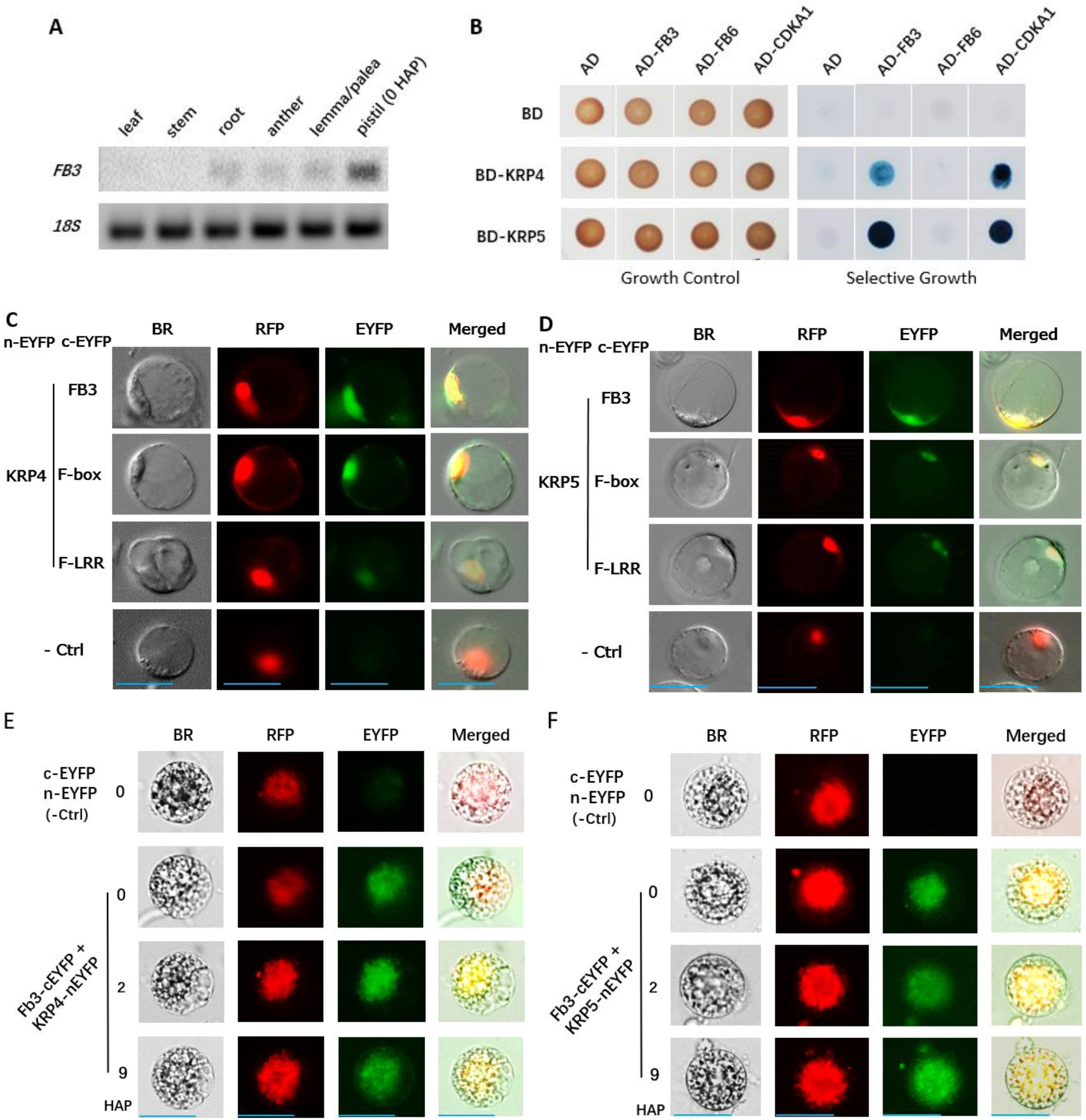
The rice FB3 protein interacts with CDK inhibitors, KRP 4 and KRP5. **A** Gel image of RT-PCR products showing expression of FB3 in six different rice organs collected from flowering rice plants (HAP, hours after pollination). 18S rRNA was used as the loading control. **B** Interactions between rice F-box proteins with rice CDK inhibitors, KRP4 or KRP5, detected in Y2H (blue spots). The preys include CDKA1 (+ control), two F-box proteins, FB3 and FB6, and the empty prey vector (AD); the baits are KRP4, KRP5, and the empty bait vector (BD). Yeast cells were grown on SD-Trp-Leu as the growth control and medium containing X-Gal with AbA as the selective growth. **C** Representative images of Bimolecular Complementation (BiFC) indicating the interaction of FB3 with KRP4 in rice leaf protoplast. The protoplasts were co-transfected with *35Spro-FB3-cEYFP, 35Spro-FB3*_*F-box*_*-cEYFP* or *35Spro-FB3*_*LRR*_*-cEYFP* with *35Spro-KRP4-nEYFP* and *35Spro-RFP (mCherry*_*NLS*_). **D** Representative images of BiFC indicating the interaction of FB3 with KRP5. Leaf protoplasts were co-transfected by *35Spro-FB3-cEYFP, 35Spro-FB3*_*F-box*_*-cEYFP* or *35Spro-FB3*_*LRR*_*-cEYFP* with *35Spro-KRP5-nEYFP and 35Spro-RFP (mCherry*_*NLS*_). In C and D, transfection with *nEYFP*+*cEYFP* was used as -Ctrl. BR indicates bright-field images. EYFP indicates the YFP signal from BiFC protein-protein interactions. RFP represents mCherry linked with Nuclear Localization Sequence (NLS) as the nuclear marker. Merged refers to the overlaid image from the other three in each transfection. Scale bars, 30 µm. **E** Representative images of BiFC to detect interactions of FB3 with KRP4 in egg cells and zygotes (2 and 9 HAP). Cells were transfected with the construct *FB3*_*pro*_*-FB3-cEYFP+Ubi*_*pro*_*-KRP4-nEYFP*. **F** Representative images of BiFC to detect interactions of FB3 with KRP5 in egg cells and zygotes (2 and 9 HAP). The cells were transfected with the construct *FB3*_*pro*_*-FB3-cEYFP+KRP5*_*pro*_*-KRP5-nEYFP*. In E and F, both *nEYFC* and *cEYFC* were used as - Ctrl; BR, cells under the bright field; mCherry_NLS_, indicates the location of the nucleus; Merged, the result of overlapping the other three images. Each combination (C-F) was repeated in three independent experiments with N = 3-6 cells per experiment. Scale bars 50 µm.

To test whether interactions between FB3 and KRP4 or KRP5 take place in rice cells, we conducted BiFC, first in rice seedling leaf protoplasts and then in isolated rice egg cells and zygotes. The clear EYFP signal in protoplasts expressing FB3-cEYFP and KRP4-nEYFP (Fig. 5C) indicates the interaction between FB3 and KRP4 and the nuclear location of the two proteins. Truncated FB3 proteins [i.e., N-terminal -100 amino acids (including the N-terminal F-box domain) or C-terminal -111 amino acids (including the C-terminal leucine-rich region domain, LRR)] also interact with KRP4 but at apparently lower level, as shown via fluorescence image quantification (Fig. 5C). Similarly, BiFC assays in protoplasts indicate a similar pattern of nuclear-localized interactions between KRP5-nEYFP and FB3-cEYFP, including the detectable interaction between KRP5-nEYFP and the truncated FB3 polypeptides (∆F-box and ∆LRR) - cEYFP (Fig. 5D).

BiFC experiments in isolated egg cells and zygotes using constructs with native promoters indicate a similar result. Detection of a nuclear-localized EYFP signal in experiments with egg and zygote cells transfected with *FB3*_*pro*_-*FB3-cEYFP* and *KRP5*_*pro*_*-KRP5-nEYFP* supports both the co-expression and protein interactions of FB3 and KRP5 (Fig. 5F). Similarly, we also detected interactions within nuclei of egg cells and zygotes transfected with *FB3*_*pro*_*-FB3-cEYFP* and *Ubi*_*pro*_*-KRP4-nEYFP* (Fig. 5E). We conclude that the rice FB3 protein interacts with the KRP4 and KRP5 proteins in the nuclei of rice egg cells and zygotes. This is consistent with a model in which FB3 regulates the inhibitory activities of KRP4 and KRP5 permitting cell cycle progression in rice early zygotes.

### FB3 degrades both KRP4 and KRP5 via the proteasome pathway

To further test the role of FB3 in controlling the activity of KRP4 and KRP5, we conducted kinase and protein degradation assays with extracts of transformed rice protoplasts, overcoming the technical challenge of very low expression of FB3 in yeast and bacteria. Transformation of rice protoplasts with GFP-tagged FB3 under the control of a β-estrogenregulated, inducible promoter system (Sreekala et al. 2005; Hirose et al. 2012) gave proteins of the expected size, 89 kDa for GFP-FB3, via western blot (Additional file 1: Fig. S3).

Applying the fluorescent ADP detection assay on mixtures of immunopurified protein complexes, we found that the inhibitory effect of KRP4 and KRP5 on CDKB1 and CDKB2 kinase activity can be reversed by FB3 (Fig. 6A). CDKB1 activity with both inhibitors presents (CDKB1 + KRP4 + KRP5) was estimated at ∼20k RFU; however, it increased by ∼2.9-fold upon the addition of purified FB3 (i.e., CDKB1 + KRP4 + KRP5 + FB3, Fig. 6A). Similarly, inhibited CDKB2 activity (CDKB2 + KRP4 + KRP5) increased by 1.7-fold, its original level, upon addition of the purified FB3 (CDKB1 + KRP4 + KRP5 + FB3). Both scenarios indicate that the two KRPs interact with FB3 and are likely degraded via FB3 mediated proteasome pathway.

**Fig. 6.**
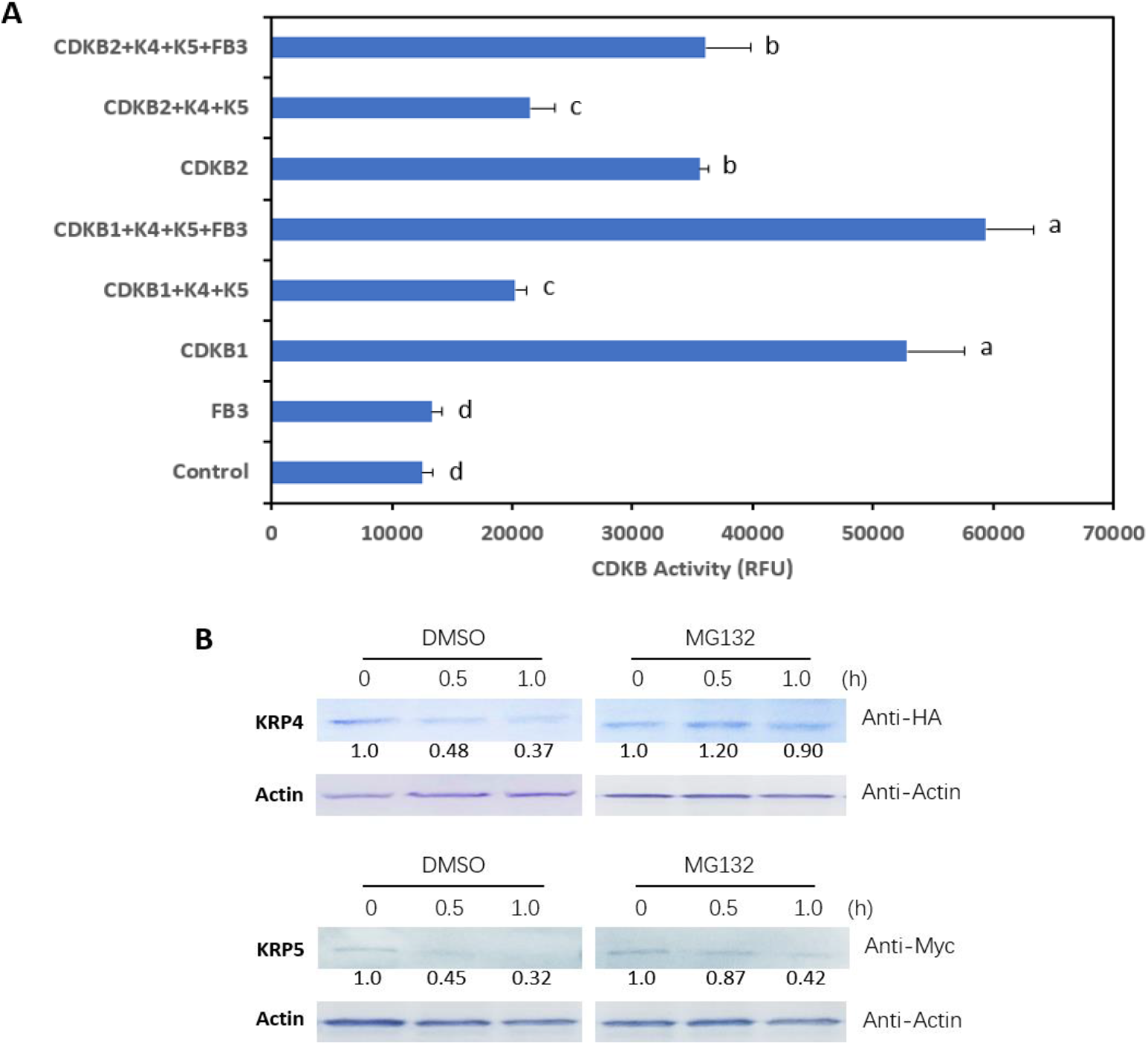
FB3 and the proteasome regulates KRP4 and KRP5. **A** Rice FB3 protein counteracts the inhibitory effect of KRP4 and KRP5 on kinase activities of CDKB1 and CDKB2. Kinase activity was measured in relative fluorescent units. The Control is with the “mock” Co-IP from extracts of yeast that express only empty vectors, i.e., the bait vector pGBKT7 and the prey vector pGADT7. Data are the mean ± standard deviation (N = 3 biological replicates). Values in A are the mean ± standard deviation, N = 3 biological repeats. Different letters indicate significantly different groups determined by one-way ANOWA analysis (*P* < 0.05). **B** The 26S proteasome-based degradation of KRP4 (the upper) or KRP5 (the bottom). Purified FB3 and KRP4 or KRP5 were mixed with rice ovary extract containing the protease inhibitor cocktail and treated with the specific proteasome inhibitor (MG132) or DMSO (control), incubated at 4°C for 0, 0.5, or 1 hour (h), and detected via western blot with anti-HA or anti-c-Myc. Quantification of band intensity is shown in numbers normalized to KRP4 or KRP5 at 0h, and Anti-actin antibody was used to show equal loading. **C** Representative images of transfected rice egg cells and zygotes with *EYFP-KRP4*. **D** Representative images of transfected rice egg cells and zygotes with *EYFP-KRP5*. **E** Representative images of transfected rice egg cell and zygotes with *EYFP-FB3*. In C to E, HAP, hour after pollination. BR, cell images in gray under bright field. RFP represents mCherry linked with Nuclear Localization Sequence (NLS) as the nuclear marker. Merged, overlapping all other images in the same transfection. Scale bars in C-E: 50 µm. **F** Protein expression profiles of KRP4, KRP5 and FB3 in rice egg cells and zygotes. Fluorescent intensity was quantified with ImageJ. Values are the mean and error bars are the standard deviation for three biological replicates (N = 3 - 6 cells in each). The * indicates a significant difference in fluorescence of KRP4 or KRP5 between 2 HAP and 9 HAP via one-way ANOVA analysis (*P* < 0.05).

In addition to interacting with each other, we obtained evidence that KRP4 and KRP5 are degraded by the 26S proteasome. After 30 to 60 min of incubation of purified KRP4 and KRP5 with purified FB3 in rice ovary extracts, western blotting with anti-Myc and anti-HA antibodies was used to detect the KRPs. Fig. 6B shows that myc-tagged KRP5 and HA-tagged KRP4 protein is degraded in rice ovary extracts (DMSO control), but that this degradation is slowed by MG132, a specific small-molecular proteasome inhibitor (Kim et al. 2008). Hence, we conclude that KRP4 and KRP5 degradations are proteasome-dependent, consistent with the involvement of an E3 ligase, such as one that includes FB3.

### Effects of the cell-cycle gene alteration on rice seed formation

To further investigate the *in vivo* functions of *KRP4, KRP5*, and *FB3* in rice embryogenesis and seed formation, we characterized five T-DNA or transposon insertion lines for the corresponding genes, consisting of two T-DNA insertion alleles for *KRP5*, a transposon insertion and a T-DNA activation tagged-allele for *KRP4*, and a transposon insertion allele for *FB3* (Fig. 7, Additional file 2: Table S3 & S4 and Additional file 1: Fig. S5-S9). Upon genotyping of the T2 generation, we identified very few homozygous insertion mutants for *krp5-1, krp5-2*, and *KRP4-D1*^*OE/+*^ and thus focused on the characterization of the heterozygotes; whereas, the *krp4-1* line was already homozygous upon the first characterization.

**Fig. 7.**
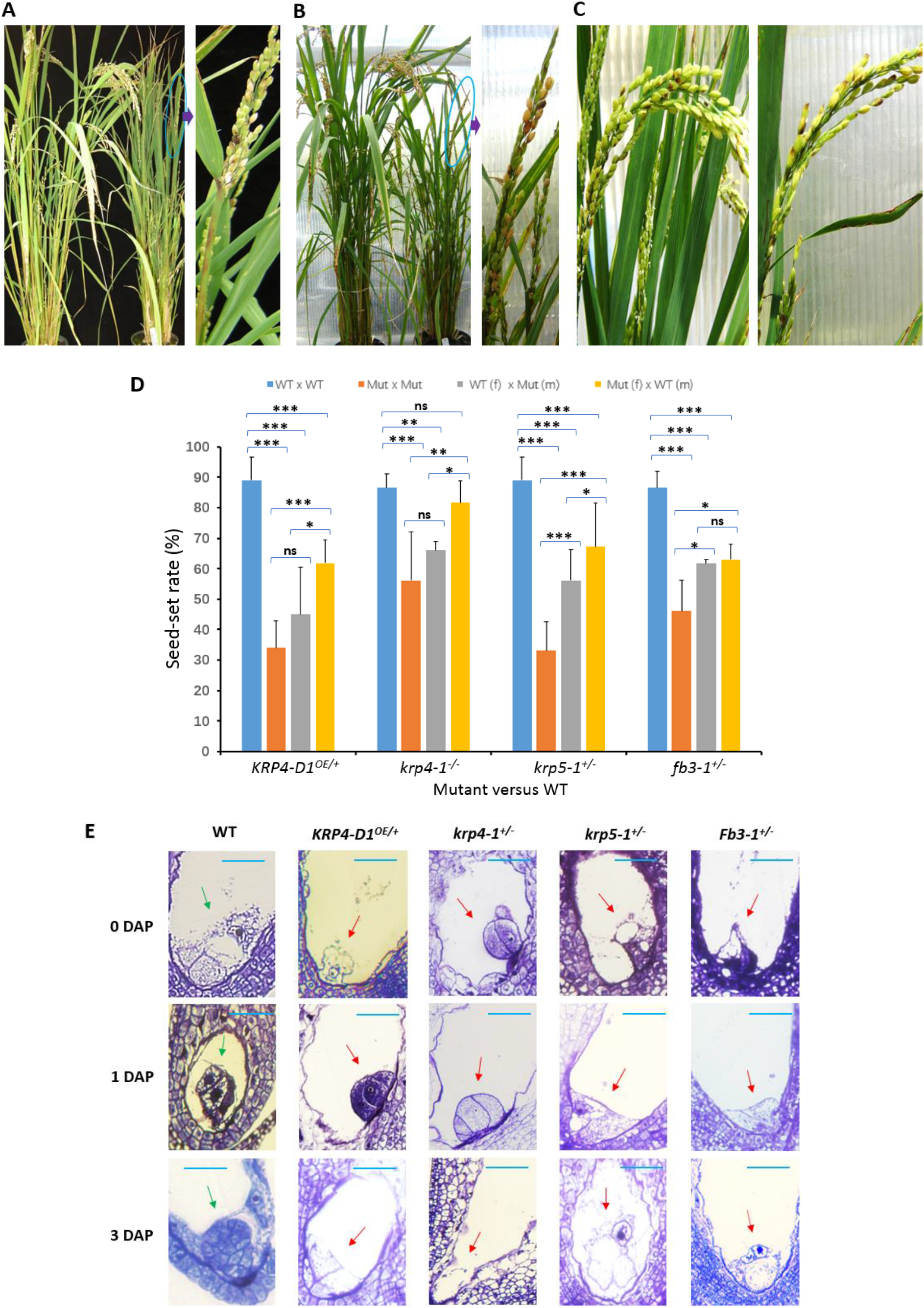
Mutations of KRP5, KRP5, and FB3 genes significantly reduce rice seed-set rate. **A** Representative WT (left) and KRP4-over-expression mutant *KRP4-D1*^*OE/* ***+***^ (right) plants. **B** Representative WT (left) and the heterozygous KRP5 knockout mutant *krp5-1*^***+/-***^ (right) plants. **C** Representative rice panicles of WT (left) and the heterozygous FB3 knockout mutant *fb3-1*^***+/-***^ (right). **D** The reduced seed-set rate in transgenic mutant lines in the T_2_ generation (n = 1000 seeds counted from 4 rice plants per line) and the partially restored seed-set rate from reciprocal pollination between rice transgenic mutants and WT in the F_o_ generation (n = 100 emasculated florets in multiple plants for each reciprocal pollination). Compared with the pollination using WT as female, pollination using WT as male significantly restored the seed-setting rates which were reduced in mutants of *krp4-1* ^***-/-***^, *KRP4-D1*^*OE/+*^, *krp5-1*^***+/-***^ *and fb3-1*^***+/-***^. Each cross is written such that the first parent listed is female (f) and the second listed parent is male (m). Columns are mean values and error bars are the standard deviation. One-way ANOVA analysis was performed to determine the significant differences among the data of different genotypes: *** indicates the difference at *P* < 0.001, ** at *P* < 0.01, and * at *P* < 0.05. **E** Micrographs of rice ovaries in WT and transgenic mutants at different stages stained with Toluidine Blue O shows abnormal cellular structure of female germ units (egg cell, synergids and central cell) and embryos in mutant ovaries (red arrows) and compares with normal, wild type (WT) in female germ units (green arrows) at different stages. The dark purple spots represent nuclei. Scale bars, 50 µm. DAP, days after pollination. WT 1 DAP is to show the two-celled pre-embryo; WT 3 DAP, a globular like embryo.

Seed-set rates were significantly reduced relative to the original parental cultivars in all mutants examined (Fig. 7), demonstrating the importance of these genes in rice gamete or zygotic development. While the seed-set rate for all parental cultivars under our growth conditions was about 90%, the seed-set rate for *fb3-1*^***+/-***^ in the T_2_ generation was reduced to 46%, which is expected for a gene required either for sperm or egg viability (0.9*50% = 45% expected). The lower seed-set rates in *krp5-1*^***+/-***^, *krp5-2*^***+/-***^, *KRP4-D1*^*OE/+*^ (33%, 28% and 34%, respectively) are consistent with haploid lethality combined with a reduction of zygote viability with unbalanced KRP abundance, i.e., halved protein abundance in the heterozygous *krp5-1*^***+/-***^, *krp5-2*^***+/***^ and increased in the *KRP4-D1*^*OE/+*^. Gametes of these lines could also have decreased viability. Consistent with KRP4 imbalance being detrimental, Yang et al (2011) also observed that overexpression of KRP4 in *indica* rice gave a low seed-set rate of 28 ± 4 (%). On the other hand, the slightly higher seed-set rate for homozygous *krp4*^***-/-***^ (56%) may be due to a partial ability of its coordinate partner, KRP5, to compensate for KRP4 deficiency in gamete cells. To address the mechanism of reduced seed-set, we examined the mutant line pollen grains and ovaries with light microscopy. No differences were apparent in *krp5-1*^***+/-***^, *KRP4-D1*^*OE/+*^, *krp4-1* ^***-/-***^ *and fb3-1*^***+/-***^ genotypes *versus* WT in FDA staining for pollen viability and sperm cell morphology via DAPI staining (Additional file 1: Fig. S12). Whereas Kim et al (2008) observed that only a single germ cell nucleus was present in the pollen of the F-box protein mutant *fbl17*, and Yang et al (2011) reported that a 33-kDa secretory protein gene (33SP) promoter driving KRP4 over expression in *indica* rice dramatically reduced pollen viability, this does not seem to be the case in the *japonica* rice mutants characterized here. Still, the sperm cells of these mutants were abnormal in function. In reciprocal pollination (Fig. 7D), we found that WT, used as either mother or father crossed with the mutants, significantly restored seed-set rates in contrast to mutant self-pollination (Fig. 7D). Moreover, compared with using mutant pollen to cross-pollinate WT, using WT as the pollen donor with mutant ovaries increased seed-set by 20% for *krp5-1*^***+/-***^, 24% for *krp4-1* ^***-/-***^ and 38% for *KRP4-D1*^*OE/+*^ (Fig. 7D), suggesting that mutation in KRP4 and KRP5 compromises rice sperm cell functions in fertilization and/or zygote formation and development.

Furthermore, examination of the morphology of the female germ unit and embryo revealed that unlike the normal cellular structure in the WT cultivars [1 egg cell, 2 synergids and 1 central cell (green arrows in Fig. 7E), in the mutants (*krp5-1*^***+/-***^, *KRP4-D1*^*OE/+*^, *krp4-1* ^***-/-***^ and *fb3-1*^***+/-***^), female germ units were collapsed at the 0 DAP and embryos were abnormally developed at 1 and 3 DAP (red arrows in Fig. 7E). Putting the above together, either functionally compromised sperm cells, structurally collapsed female germ unit cells or irregularly developed embryos may negatively impact the cell cycle during embryogenesis thus significantly lower rice see-set rate.

To follow up, we conducted additional cross-pollination between two rice mutant lines for two purposes, examining the phenotype of a *krp4krp5* double mutant and *krp4-1* complementation. A double mutant line, *krp4-1*^***-/-***^*krp5-1*^***+/-***^, would have allowed further exploration of the functional redundancy of the two inhibitors during the rice zygotic cell cycle. However, though all of the few seeds obtained germinated, none of the double mutant plants survived past early seedling stage (Table 1). The low seed-set is consistent with a crucial role of coordination between KRP4 and KRP5 in rice zygotic development and grain formation. The failure of the obtained seedlings to develop suggests that the KRP4-KRP5 complex may have other functions in vegetative development not revealed by the study of hemizygous mutants. The other objective was to complement the mutant *krp4-1*^***-/-***^ with the *KRP4-D1*^*OE/+*^ activation tagged line. In this case, cross-pollination was successful though only 3 filled seeds were harvested. We were able to grow these to maturity and found that all the progeny carry KRP4 OE allele by genotyping and their seed-set rate was restored to 74% (Table 1), providing evidence for partial complementation of the KRP4 abundance inbalance in the progeny. Since the mutant krp4-/- plants have a partial seed set of ∼ 50% (Fig. 7D), only ∼ 24% was rescued from the remaining 50% (non-seed set). Therefore, the rescue frequency is actually 50% (ratio of 1:1), as expected from segregation of the complementing *KRP4-D1*^*OE*^ transgene.

**Table 1.** Results of cross-pollination between selected rice mutant lines

| Purpose | Cross<sup>a</sup> | Hand-Pollinated-Florets (#) | Seeds harvested (#) | Progeny survival (%) | Seed-set (%) |
| --- | --- | --- | --- | --- | --- |
| Control | WT (Nip) x WT (DJ) | 100 | 93 filled | 95 | 90 |
| Obtain double mutant | *krp4-1<sup>-/-</sup>* x *krp5-1<sup>+/-</sup>* | 200 | 1 filled + 5 half-filled | 0 | 0 |
| *KRP4* complementation | *krp4-1<sup>-/-</sup>* x *KRP4-D1<sup>OE/+</sup>* | 100 | 3 filled + 5 half-filled | 38<sup>b</sup> | 74 |

<sup>a</sup> *Nip*, Nipponbare; *DJ*, Dongjing
<sup>b</sup> The survived progeny was genotyped: all carried the OE allele.

## Discussion

### The relationship of KRP4 and KRP5 in plant zygotic cell cycle control and their degradation by rice F-box proteins suggests the important role of the cell cycle proteins in the formation of zygotes and seeds

Based on our research, a model of rice zygotic cell cycle control can be described (Fig. 8). The core complex consists of four members: CDKB1, CYCD5, KRP4 and KRP5. This model shows that the activity of CDKB1 is CYCD5-dependent but inhibited by KRP4 and KRP5 prior to zygotic division. KRP4 and KRP5 physically interact to cooperate in inhibiting CDKB1 activity.

**Fig. 8.**
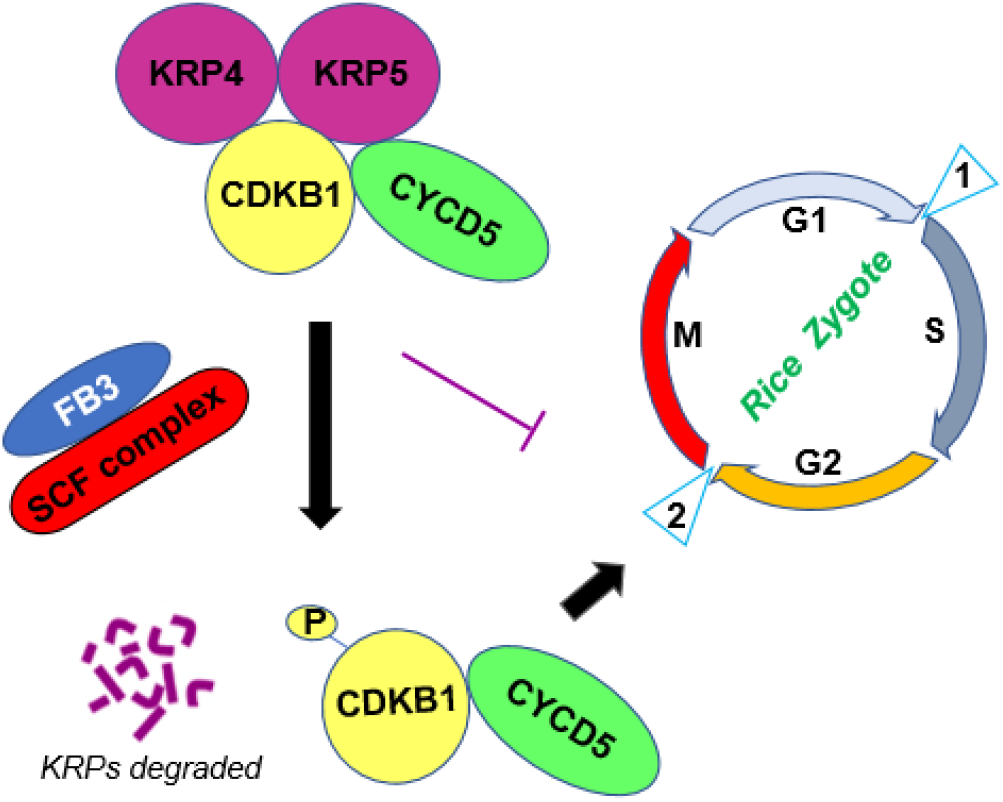
Model of the core complex for rice zygotic cell cycle control at checkpoint 2, Gap 2 to Mitosis (G2-M). CYCD5 dependent CDKB1 is coordinately inhibited by KRP4 and KRP5, and the two inhibitors are degraded via the FB3-mediated proteasome pathway, which then permits the cell cycle to proceed.

The advantages of such a relationship for cell cycle control might be improved stabililty and redundancy if one of the two KRPs is dysfunctional. KRP4-KRP5 complex formation is supported by multiple lines of evidence, including yeast two hybridization, co-immunoprecipitation, yeast growth, *in vitro* CDK activity, BiFC in rice zygotes and mutant analysis. To our knowledge, this is the first report of a coordinate relationship of CDK inhibitors. Based on coexpression evidence, it seems likely that coordinated function is a universal mechanism for KRP-mediated cell cycle control in plants. For example, both AtKRP6 and AtKRP7 are expressed during spermatogenesis of Arabidopsis (Kim et al. 2008), and both KRP1 and KRP2 are preferentially expressed in rice sperm cells as (Fig. 1A).

As cell cycle inhibitors, if KRP4 or KRP5 hyper accumulated, zygotes could be prevented from entering G2 or Mitosis, which is supported by our results with *KRP5* overexpression. On the other hand, if KRP4 or KRP5 abundance is reduced, zygotes might have been expected to prematurely advance to G2 or Mitosis, which could result in bigger seeds. Unfortunately, this yield advantage is not observed (Fig. 7). Instead, our results suggest that an abundance balance between the KRPs and other cell cycle proteins is crucial for the development of gametes, zygotes, embryo, and endosperm.

FBPs compose a superfamily in plants. Through the proteasome pathway, they are involved in many different processes in plant growth and development including cell cycle control, signal transduction, metabolic regulation, floral organogenesis, and senescence. Another important reproductive event, plant sperm cell formation is under the control of two cell cycle inhibitors, AtKRP5 and AtKRP6, which are degraded by the FBL17-associated SCF complex (Kim et al. 2008). Here, we identify FB3 as a novel rice F-box protein preferentially expressed in rice egg cells and zygotes and with a role in degrading both KRP4 and KRP5 via the 26S proteasome pathway. Experimental evidence for this function consists of the protein degradation assay, reversal of KRP-mediated inhibition of CDKB activity, and interaction of FB3 with KRP4 and KRP5 as indicated in Y2H, BiFC, and cellular localization in rice zygotes at different stages. Our results demonstrate that FB3 is a regulator of the two KRP inhibitors for rice zygotic cell cycle control.

We also confirmed that mutations in KRP4, KRP5 and FB3 result in the compromised function of sperm cells and abnormal structure of the female germ unit and embryo in the mutants, and hence leading to significantly reduced seed-sets. These results support that balanced ratio of KRP4, KRP5 and FB3 proteins are important for zygotic development. Since the zygote is formed from the combination of one male and one female gamete, defective gametes fail to contribute the three proteins to the zygote. Imbalances of these proteins in the zygote appear to frequently lead to the failure of embryogenesis and seed formation (Fig. 7), though more evidence to understand whether these effects are quantitative or qualitative is necessary. Still, our data also provides insight into zygotic cell cycle control in plants.

In addition, RNAseq data shows that CDKBs (Additional file 1: Fig. S11A) are present in rice egg cells but preferentially expressed in rice zygotes (9 and 14 HAP), and CYCD5 (Additional file 1: Fig. S11B) is preferentially expressed in rice egg cells and zygotes, supporting them as the components in our cell cycle model; meanwhile, some other transcripts like CDKC1;2 and CYCT1;4 are also abundant in rice egg cell and zygotes (Additional file 1: Fig. S11). Further study is required to reveal their functions.

## Conclusion

In this STUDY, we characterized a novel core complex of cell cycle proteins for rice zygotic mitosis. The complex consists of Cyclin D5, Cyclin-Dependent Protein Kinase B1, two coordinately Kip-Related Proteins, KRP4 and KRP5, and the F-box protein FB3, which regulates the two KRPs. The genetic data of whole plant and microscopy data of rice mutant ovary also support the role of this complex in rice egg cells and initiation of embryogenesis.

## Materials and Methods

### Plant material

Rice seeds were germinated in the dark at room temperature for 5 days. Seedlings were transplanted to soil and maintained in a greenhouse (27°C during daytime, 12 hours, and 25°C for nighttime, 12 hours) until blooming. The daytime lighting was automatically controlled to 500 μmol m^-2^s^-1^. Plants were irrigated with deionized water and fertilized twice weekly by filling the headspace in pots with 300-350 ppm nitrogen (Jack’s 20-10-20). Mutant lines (Supplementary Table S4) were identified from the Rice Functional Genomic Express Database (http://signal.salk.edu/cgi-bin/RiceGE) and obtained from South Korea (Jeong et al. 2002 and 2006) and UC-Davis (Kumar et al. 2005).

### RNA isolation and RT-qPCR

For RNA extraction, samples (100 mg each) were collected from 8-week-old plants. Fresh florets were collected and immersed in 0.3 M mannitol for dissecting anthers, lemma/palea, and pistil followed by snap freezing in liquid nitrogen. Total RNA was extracted using Qiagen RNAeasy Plant Mini Kit. Reverse transcription (RT) was performed with RevertAid First Strand cDNA Synthesis Kit (Thermo Scientific). RT-PCR was conducted on BioRad C1000 Touch Thermal Cycler system with Phusion High Fidelity (Thermo Scientific). For RT-qPCR, PowerUp™SYBR™ Green Master Mix Kit (Thermo Scientific) was employed and 18S rRNA gene was used as the internal control based on consistent expression in eight different rice samples of all three biological replicates. The relative transcript abundances from RT-qPCR were calculated using the 2^− ΔΔCt^ method (Schmittgen and Livak 2008). Primers are listed in Additional file 2: Table S5.

### Yeast two hybridization (Y2H) assays

Y2H was conducted using Matchmaker Gold Yeast Two-Hybrid System (Clontech). The coding sequences (CDS) of rice *KRP1, KRP4, KRP5* and 5’-most-315 bp of *KRP4* (encoding KRP4m, which has 90 aa deleted from C-terminus of KRP4) and 5’-most 530 bp of *KRP5* (encoding KRP5m, which has 48 aa deleted from the C-terminus of KRP5) were amplified and cloned into vector pGBKT7 as bait constructs; CDSs of CDKs, CYCs and F-box genes (Additional file 2: Table S2) were cloned into pGADT7 as prey constructs. The transformed or hybrid cells were cultured on the selective medium of DDO (SD/-Trp-Leu) or QDO (SD/-Trp-Leu-His-Ade) both containing X-ɑ-gal and Aureobasidin A (AbA) at 30°C for 3 days. Protein-protein interactions between baits and preys are indicted by blue stained patches of yeast growth.

### Isolation of yeast nuclei for nuclear protein KRP4 and KRP5 assays

The isolation of yeast nuclei was as previously described (Reese et al., 2008) with the following modifications. The yeast cells containing c-Myc or HA-tagged constructs (c-Myc-KRP5, cMyc-KRP4, HA-CDKB1, and HA-CDKB2) were cultured on -Leu-Trp plate at 30°C for 3 days, then cultured in -Leu-Trp media (up to 200 ml). The collected yeast cells were treated with cell-wall enzyme solution containing 2-3 mg Zymolyase 100T (USB) (30°C, 2 hs). Spheroplasts were pelleted (4500 ***g***, 4°C, 5 minutes) in 10 ml -Leu-Trp/S, washed twice in 25 ml cold -Leu-Trp/S and once in 20 ml 1M Sorbitol (ice cold), then homogenized on ice with Nuclei buffer (30 mM Hepes pH7.6, 0.05 mM CaCl_2_, 5 mM MgSO_4_.7H_2_O, 1 mM EDTA, 10% Glycerol, 0.5% NP40, 7.2 mM Spermidine Hexahydrate with protease inhibitors (Sigma), PMSF (1 mM) and ß-me (2 mM). After removing cellular debris, nuclei were collected from the supernatant (14000 ***g***, 4°C, and 10 min) and stored at -20°C.

### Co-immunoprecipitation (Co-IP) assay

Pierce™ Co-Immunoprecipitation Kit (Thermo Scientific™) was used. Frozen yeast nuclei were suspended in Lysis Buffer containing 25 mM Tris (pH7.5), 150 mM NaCl, 1 mM EDTA, 1% NP40, 5% glycerol 10 mM NaF, 1 mM Na_3_VO_4_ and freshly added with protease inhibitor cocktail (Sigma) plus PMSF (1 mM) and incubated on ice for 10-15 minutes with agitation. The cell debris was removed by centrifugation (13,500 ***g***, 4°C, and 8 min), and resulting supernatant was saved on ice: 1 µl was used to determine protein concentration and the rest was precleared with 40 µl control agarose beads. About 25 µg of anti-HA (GenScript) or anti-c-Myc mAb (GenScript) were coupled with 50 µl beads and incubated with precleared proteins at 4°C for overnight with gentle mixing. Finally, proteins were eluted with elution buffer (pH 2.8) containing primary amine.

### Western blotting

The eluted protein (∼10 µg) was separated within 12% SDS - PAGE and transferred to PVDF membrane (Bio-Rad). The membrane was blocked for 1 hour, then incubated with primary antibody (Anti-HA or anti-c-Myc, mouse mAb) at 4°C for overnight, washed for three times and incubated with horseradish peroxidase (HRP) - conjugated secondary antibodies (goat anti-mouse, Invitrogen) for 1 hour. After washing the membrane was detected with One Step Ultra TMB-blotting Solution (Thermo Fisher) or SuperSignal™ West Pico PLUS Chemiluminescent Substrate (Thermo Scientific).

### CDK activity assay

Kinase activity was measured using ADPsensor™ Universal Kinase Activity Assay Kit (BioVision). HA-tagged CDKB1, CDKB2 and CYCD5 and c-Myc-tagged KRP4 and KRP5 were purified from yeast nuclei, and FB3-GFP from suspension-cultured rice protoplasts, by immunoprecipitation in antibody (each 20 µg) immobilized agarose beads (each 50 µl) from Pierce™ Co-Immunoprecipitation Kit (Thermo Scientific™). For each assay, 1 µg of purified protein was used; for 1 µg of FB3, 6 µl of rice ovary extract (containing 26S proteasome) was also added. The fluorescence was scanned at Ex/Em = 535/587 nm in a multimode plate reader (BioTek, Synergy™ HT).

### Rice cell isolation and transfection for BiFC and cellular localization

Isolation and transfection of rice leaf protoplasts were conducted as previously described (Wang et al. 2013). Isolation of rice egg cells and zygotes was conducted as previously described (Li et al. 2019, 2020 and 2022). Transfection of isolated rice egg cells and zygotes was based on previous reports (Koiso et al. 2017; Toda et al. 2019) with modifications. First, several isolated cells were transferred on to a micro slide (within a moisture chamber) in 8 µl MMG (4 mM Mes-KOH, pH5.7, 15 mM MgCl_2_ and 0.4 M mannitol), added with 1-2 µg plasmid DNA and 10 µl of 30% PEG with 100 mM CaCl_2_ in 0.4 M mannitol, followed by gently mixing with a micro-pipet. After incubation for 10 min, the cells were carefully washed 3 times with fresh MMG (capillary replacement of ∼ 10 µl each time), followed by 3 more washes in modified W5 solution (W5: 154 mM NaCl, 125 mM CaCl_2_, 5 mM KCl, 2 mM MES; plus 500 mM Glucose). Then, the cells were cultured with ∼ 10 µl of the modified W5 solution containing Ampicillin (100 µg/ml) at 25°C overnight in a dark moist chamber.

### Constructs for Bimolecular Fluorescent Complementation (BiFC)

The CDS of *KRP4* and *KRP5* were cloned into pE2913 (Lee and Gelvin 2014) for construct *KRP4-nEYFP or KRP5-nEYFP*, and the CDS of *CDKB1, KRP4*, and *FB3* into pE2914 (Lee and Gelvin 2014) for *CDKB1-cEYFP, KRP4-cEYFP* or *FB3-cEYFP*; the rice promoter *KRP5* (*KRP5*_*pro*_), maize promoter *Ubiquitin* (*Ubiq*_*pro*_), and rice promoter *CDKB1* (*CDKB1*_*pro*_) were added to 5’ termini of the construct *KRP5-nEYFP, KRP4-nEYFP, KRP4-cEYFP*, and *CDKB1-cEYFP*, respectively. To test the interaction of CDKB1 and KRP4 or KRP5, the construct *CDKB1*_*pro*_*-CDKB1-cEYFP* was linked to *Ubi*_*pro*_*-KRP4-nEYFP* and *KRP5*_*pro*_*-KRP5-nEYFP*, separately, using Gibson Assembly kit (NEB). To test the interaction of KRP4 and KRP5, the construct *Ubi*_*pro*_*-KRP4-cEYFP* was linked to *KRP5*_*pro*_*-KRP5-nEYFP* via Gibson Assembly. Primers are listed in Additional file 2: Table S5.

### Suspension culture to express FB3 protein in rice protoplasts

The ß-estradiol inducible expression system was used to express FB3 in protoplasts. For this, the CDS of *FB3* was linked to GFP in pUH-GFP2 (Sreekala et al. 2005) and transfected into leaf protoplasts (Wang et al. 2013). The transfected cells were cultured for 6-9 days (26°C, dark with shaking at 80 rpm) in Chu’s medium (3.99 g/l, Caisson Cat# CHP03) with Hygromycin (35 mg/l), Ampicillin (100 mg/l) and ß-estradiol (inducer, 20 µM) plus supplementals (Chen et al. 2017; Chen et al. 2006; Kim et al. 2008): 2,4-D (2 mg/l), casamino acids (300 mg/l), L-glutamine (500∼1000 mg/l), L-proline (1000 mg/l), pyridoxine (1.0 mg/l), kinetin (0.2 mg/l), biotin (0.01 mg/l) and retinol (0.01 mg/l) and Sucrose (30 g/l.). Fresh sterile Chu’s medium was used to expand the culture (1:3 in volume) every 3 days.

### Protein extraction from rice ovary (for 26S proteasomes with SCF complexes)

About 200 ovaries were homogenized on ice in 0.5 ml protein degradation buffer [25 mM Tris pH7.4, 150 mM NaCl, 1 mM EDTA, 1% NP40, and 5% glycerol with freshly added plant protease inhibitor cocktail (Sigma) and 1 mM PMSF]. Cellular debris was pelleted at 3000 g, 4°C for 5 min, and the supernatant was aliquoted and stored at -20°C.

### Protein degradation assay

The purified proteins of KRP4 and KRP5 (tagged with c-Myc) and FB3 protein (tagged with GFP) were added to a degradation assay based on Kim et al (2008). Briefly, ovary extract (100 µl) was pretreated with the proteasome inhibitor, MG132 (10 µM), to slow protein degradation. Extracts were then mixed with purified KRP4 or KRP5 and FB3 proteins (1 µg each); in the control, MG132 was replaced by DMSO (1%) for normal protein degradation. The two mixtures were incubated at 4°C and sampled (16 µl) every 30 minutes. KRP4 or KRP5 was detected with anti-HA or anti-Myc antibody in western blot, and Actin was used as the loading control.

### Genomic DNA isolation and genotyping

Leaf tissue was collected from young plants (∼200 mg, 2-4 weeks) and frozen in liquid nitrogen followed by grinding in Geno Grinder (SPEX SamplePrep 2010) at 1450 rpm for 60 seconds. The sample was mixed with 200 µl of Extraction Buffer (100 mM Tris-Cl, pH 8.0, 1 M KCl and 10 mM EDTA), heated (60°C, 30 min) and extracted with 600 µl of dH_2_O (14,000 ***g***, 10 min). Supernatant (1 µl) was used for genotyping with primers (Additional file 2: Table S6).

### Seed-setting assay

Rice seeds (1000) were harvested from 3-5 plants of each wildtype or mutant line and counted for the number of filled or aborted seeds. The seed-set rate was calculated as (number of filled seeds / total number of seeds) x 100 %.

### Cross-pollination

Crossing was conducted following the McCouch Lab procedure (2016). To ensure pollination, the glassine bags covering panicles for pollination were tapped every half hour from 11 am to 2 pm for 3 days. The seeds were counted in 1 week and harvested in 4 weeks.

### Light microscopy of rice ovary

Based on previous description (Skepper and Powell 2008). 3-5 rice ovaries were dissected from mature florets without pollination (day 0) or 1 or 3 days after pollination, and fixed with 2% paraformaldehyde (PFA) and 2% glutaraldehyde (GA) in 50 mM PIPES (pH 6.8), with 1 mM MgSO_4_, 5 mM EGTA. After dehydration, samples were infiltrated with LR White resin (Sigma), then transferred to gelatin capsules containing fresh resin with the appropriate orientation and polymerized at 60°C for 24-48 hours in a fume hood. After sectioning with a microtome, sections (2 μm) were stained with Toluidine Blue O (0.5% in 2% Sodium Borate) for microscopy.

## Acknowledgements

We thank Dr. Christina Bourne for valuable advices and practical help for project progression and manuscript preparation; Dr. Anne Dunn for providing instruments, chemicals, and financial support for western blotting analysis; Mr. Lynn Nichols for helping manage rice plant growth; Mr. Joshua Chesnut for demonstrating manual isolation of rice egg cells; Professor Chua Nam Hai (Rockefeller University) and Dr. Yingzhong Chao for generously providing the vectors pUH-GFP2 and pER8; Dr. Paul Sims for offering the fluorescent plate reader for measuring kinase activity; Dr. Ben Smith and Dr. David Thomas for instructions in epi-fluorescence microscopy; Dr. Ann West, Dr. Sharon Kessller and Dr. Daniel Jones for constructive suggestions at the initiation of this work; Dr. Dirk Anderson and Dr. You Zhou for light microscopy of rice ovary; and the University of Oklahoma for Robberson Scholarship.

## Author Contributions

H.X. designed and performed all experiments, analyzed data, and drafted the manuscript. L.B. organized and managed analysis of multiple rice mutants, assisted with qPCR performance and participated in manuscript preparation. M.L. provided vectors and participated in molecular cloning and manuscript preparation. V.S. coordinated and supported the entire research project and provided rice mutant lines, RNA-seq data and instructive suggestions. H.F. managed rice plant growth and participated in the routine cell isolation. S.R. supervised and supported the entire research project, managed rice cell isolation, and participated in manuscript preparation.

## Data availability

All data supporting this work are available from the published online as stated within the main text and supplementary information.

## Ethics Approval and Consent to Participate

Not applicable.

## References

Ajadi AA, Tong X, Wang H, Zhao J, Tang L, Li Z, Liu X, Shu Y, Li S, Wang S, Liu W, Tajo SM, Zhang J, Wang Y (2020) Cyclin-dependent kinase inhibitors KRP1 and KRP2 are involved in grain filling and seed germination in rice (Oryza sativa L.). International Journal of Molecular Sciences 21:1–16.

Andersen SU, Buechel S, Zhao Z, Ljung K, Novák O, Busch W, Schuster C, Lohmann JU (2008) Requirement of B2-typecyclin-dependent kinases for meristem integrity in Arabidopsis thaliana. Plant Cell 20:88–100.

Anderson SN, Johnson CS, Chesnut J, Jones DS, Khanday I, Woodhouse M, Li C, Conrad LJ, Russell SD, Sundaresan, V (2017) The zygotic transition is initiated in unicellular plant zygotes with asymmetric activation of parental genomes. Dev Cell 43:349–358.

Anderson SN, Johnson CS, Jones DS, Conrad LJ, Gou X, Russell SD, Sundaresan V (2013) Transcriptomes of isolated Oryza sativa gametes characterized by deep sequencing: evidence for distinct sex-dependent chromatin and epigenetic states before fertilization. Plant J 76:729–741.

Atkins KC, Cross FR (2018) Interregulation of CDKA/CDK1 and the Plant-Specific Cyclin-Dependent Kinase CDKB in Control of the Chlamydomonas Cell Cycle. Plant Cell 30: 429–446.

Avella M, Xiong B, Dean J. 2013. The molecular basis of gamete recognition in mice and humans. Mol. Hum. Reprod 19:279–289.

Bai C, Sen P, Hofmann K, Ma M, Goebl M, Harper JW, Elle SJ (1996) SKP1 connects cell cycle regulators to the ubiquitin proteolysis machinery through a novel motif, the F-box. Cell 86:263–274.

Barrôco RM, Peres A, Droual AM, De Veylder L, Nguyen le SL, De Wolf J, Mironov V, Peerbolte R, Beemster GT, Inzé D, Broekaert WF, Frankard V (2006) The cyclin-dependent kinase inhibitor Oryza; KRP1 plays an important role in seed development of rice. Plant Physiol 42: 1053–1064.

Boudolf V, Vlieghe K, Beemster GTS, Magyar Z, Acosta JAT, Maes S, Schueren EVD, Inzé D, De Veylder L (2004) The plant-specific cyclin-dependent kinaseCDKB1;1 and transcription factor E2Fa-DPa control the balance of mitotically dividing and endoreduplicating cells in Arabidopsis. Plant Cell 16:2683–2692.

Chamberlin MA, Lawi SJ (2017) Development and observation of mature megagametophyte cell-specific fluorescent markers. In: Schmidt A (ed) Plant Germline Development, Methods and Protocols. Humana, New York, pp 55–65.

Cheng Y, Cao L, Wang S, Li Y, Shi X, Liu H, Li L, Zhang Z, Fowke LC, Wang H, Zhou Y (2013) Downregulation of multiple CDK inhibitor ICK/KRP genes upregulates the E2F pathway and increases cell proliferation, and organ and seed sizes in Arabidopsis. Plant J 75:642–655.

Chen S, Tao L, Zeng L, Vega-Sanchez ME, Umemura K, Wang GL (2006) A highly efficient transient protoplast system for analyzing defense gene expression and protein-protein interactions in rice. Mol Plant Pathol 7:417–427.

Chen Z, Cheng Q, Hu C, Guo X, Chen Z, Lin Y, Hu T, Bellizzi M, Lu G, Wang GL, Wang Z, Chen S, Wang F (2017) A chemical-induced, seed-soaking activation procedure for regulated gene expression in rice. Front Plant Sci 8:1–13.

Cyprys P, Lindemeier M, Sprunck S (2019) Gamete fusion is facilitated by two sperm cell expressed DUF679 membrane proteins. Nature Plants 5:253–257.

Dante RA, Larkins BA, Sabelli PA (2014) Cell cycle control and seed development. Frontiers in Plant Science 493:1–14.

De Veylder L, Beeckman T, Beemster GT, Krols L, Terras F, Landrieu I, van der Schueren E, Maes S, Naudts M, Inzé D (2001) Functional analysis of cyclin-dependent kinase inhibitors of Arabidopsis. Plant Cell 13:1653–1668.

De Veylder L, Beeckman T, Inzé, D (2007) The ins and outs of the plant cell cycle. Nature Reviews Molecular Cell Biology 8:655–665.

Dresselhaus T, Sprunck S, Wessel GM. 2016. Fertilization Mechanisms in Flowering Plants. Current Biology 26(3):R125–139.

Dudits D, Cserháti M, Miskolczi P, Horváth GV (2007) Cell Cycle Control and Plant Development: The growing family of plant Cyclin-Dependent Kinases with multiple functions in cellular and developmental regulation. Annual Plant Reviews 32:1–30.

Godfray HCJ, Beddington JR, Crute IR, Haddad L, Lawrence D, Muir JF (2010) Food security: the challenge of feeding 9 billion people. Science 327:812–818.

Guo J, Song J, Wang F, Zhang XS (2007) Genome-wide identification and expression analysis of rice cell cycle genes. Plant Mol Biol 64:349–60.

Hirose T, Mizutani R, Mitsui T, Terao TA (2012) Chemically inducible gene expression system and its application to inducible gene suppression in rice. Plant Prod Sci 15:73–78.

Hu SY (1998) Centenary on SG Nawaschin’s discovery of double fertilization: retrospects and prospects. Acta Bot Sin 40:1–13.

Hu X, Cheng X, Jiang H, Zhu S, Cheng B, Xiang Y (2010) Genome-wide analysis of cyclins in maize (Zea mays). Genet Mol Res 9:1490–1503.

Jeong DH, An S, Kang HG, Moon S, Han JJ, Park S, Lee HS, An K, An G (2002) T-DNA insertional mutagenesis for activation tagging in rice. Plant Physiol 130:1636–1644.

Jeong DH, An S, Park, S, Kang HG, Park GG, Kim SR, Sim J, Kim YO, Kim MK, Kim SR, Kim J, Shin M, Jung M, An G (2006) Generation of a flanking sequence-tag database for activation tagging lines in japonica rice. Plant J 45:123–132.

Khanday I, Sundaresan V (2021) Plant zygote development: recent insights and applications to clonal seeds. Current Opinion in Plant Biology 13:1–10.

Kim HJ, Oh SA, Brownfield L, Hong SH, Ryu H, Hwang I, Twell D, Nam HG (2008) Control of plant germline proliferation by SCFFBL 17 degradation of cell cycle inhibitors. Nature 455:1134–1137.

Kim TG, Baek MY, Lee EK, Kwon TH, Yang MS (2008) Expression of human growth hormone in transgenic rice cell suspension culture. Plant Cell Rep 27:885–891.

Kipreos ET. Pagano M (2000) The F-box protein family. Genome Biol 1:1–7.

Koiso N, Toda E, Ichikawa M, Kato N, Okamoto T (2017) Development of gene expression system in egg cells and zygotes isolated in rice and maize. Plant Direct 1:1–10.

Kumar CS, Wing RA, Sundaresan V (2005) Efficient insertional mutagenesis in rice using the maize En/Spm elements. Plant J 44:879–892.

La H, Li J, Ji Z, Cheng Y, Li X, Jiang S, Venkatesh PN, Ramachandran S (2006) Genome-wide analysis of cyclin family in rice (Oryza Sativa L.). Mol Genet Genomics 275:374–386.

Lee L, Gelvin SB (2014) Bimolecular Fluorescence Complementation for imaging protein interactions in plant hosts of microbial pathogens. In: Vergunst AC, O’Callaghan d (eds) Host-Bacteria Interactions, Methods and Protocols. Humana, New York, pp 185–208.

Leene JV, Boruc J, De Jaeger G, Russinova E, De Veylder L (2011) A kaleidoscopic view of the Arabidopsis core cell cycle interactome. Trends in Plant Science 16:141–150.

Li C, Gent JI, Xu H, Fu H, Russell SD and Sundaresan V (2022) Resetting of 24-nt siRNA landscape is initiated in the unicellular zygote in rice. Genome Res 32:309–323.

Li C, Xu H, Fu FF, Russell SD, Sundaresan V, Gent JI (2020) Genome-wide redistribution of 24-nt siRNAs in rice gametes. Genome Res 30:173–184.

Li C, Xu H, Russell SD, Sundaresan V (2019) Step-by-step protocols for rice gamete isolation. Plant Reprod 32:5–13.

Lin, HY, Chen, JC, Wei MJ, Lien YC, Li HH, Ko SS, Liu ZH, Fang SC (2014) Genome-wide annotation, expression profiling, and protein interaction studies of the core cell-cycle genes in Phalaenopsis Aphrodite. Plant Mol Biol 84:203–26.

McCouch S (2016) How to cross-pollination rice in the greenhouse. http://ricelab.plbr.cornell.edu/cross_pollinating_rice. Accessed January 2022.

Ma Z, Wu Y, Jin J, Yan J, Kuang S, Zhou M, Zhang Y, Guo AY (2013) Phylogenetic analysis reveals the evolution and diversification of cyclins in eukaryotes. Mol Phylogenet Evol 66:1002–1010.

Malik Z, Gul A, Amir R, Munir F, Babar MM, Bakhtiar SM, Hayat MQ, Paracha RZ, Khalid Z, Alipour H (2020) Classification and Computational Analysis of Arabidopsis thaliana Sperm Cell-Specific F-Box Protein Gene 3p. AtFBP113. Front Genet 11:1–17.

Mizutani M, Naganuma T, Tsutsumi KN, Saitoh Y (2010) The syncytium-specific expression of the Orysa; KRP3 CDK inhibitor: implication of its involvement in the cell cycle control in the rice (Oryza sativa L.) syncytial endosperm. J Exp Bot 61:791–798.

Mori T, Igawa T, Tamiya T, Miyagishima S, Berger F (2014) Gamete Attachment Requires GEX2 for Successful Fertilization in Arabidopsis. Current Biology 24:170–175.

Mori T, Kuroiwa H, Higashiyama T, Kuroiwa T (2006) GENERATIVE CELL SPECIFIC 1 is essential for angiosperm fertilization. Nat Cell Biol 8:64–71.

Otto T, Sicinski P (2017) Cell cycle proteins as promising targets in cancer therapy. Nat Rev Cancer 17: 93–115.

Pagano M, Tam SW, Theodoras AM, Beer-Romero P, Sal GD, Chau V, Yew PR, Draetta GF, Rolfe M (1995) Role of the ubiquitin-proteasome pathway in regulating abundance of the cyclin-dependent kinase inhibitor p27. Science 269:682–685.

Pedroza-Garcia JA, Domenichini S, Raynaud, C (2016) Plant cell cycle transitions. In: Rose RJ (ed) Molecular Cell Biology of the Growth and Differentiation of Plant Cells. CRC, London, pp 1–21.

Reese JC, Zhang H, Zhang Z (2008) Isolation of Highly Purified Yeast Nuclei for Nuclease Mapping of Chromatin Structure. Methods in Molecular Biology 463:43–53.

Russell SD (1992) Double fertilization. International Review of Cytology 140:357–388.

Schmittgen TD, Livak, KJ (2008). Analyzing real-time PCR data by the comparative C-T method. Nat Protoc 3:1101–1108.

Schulman BA, Carrano AC, Jeffrey PD, Bowen Z, Kinnucan ER, Finnin MS, Elledge S, Harper JW, Pagano M, Pavletich NP (2000) Insights into SCF ubiquitin ligases from the structure of the Skp1–Skp2 complex. Nature 408:381–386.

Skepper JN, Powell JM (2008) Ultrastructural Immunochemistry: Immunostaining of London Resin (LR) White section for TEM. Cold Spring Harbor Protocols 3:1–5.

Sprunck S (2020) Twice the fun, double the trouble: gamete interactions in flowering plants. Opinion in Plant Biology 53:106–116.

Sprunck S, Rademacher S, Vogler F, Gheyselinck J, Grossniklaus U, Dresselhaus T (2012) Egg cell secreted EC1 triggers sperm cell activation during double fertilization. Science 338:1093–1097.

Sreekala C, Wu L, Gu K, Wang D, Tian D, Yin Z (2005) Excision of a selectable marker in transgenic rice (Oryza sativa L.) using a chemically regulated Cre/loxP system. Plant Cell Reports 24:86–94.

Stals H, Inze D (2001) When plant cells decide to divide. Trends in Plant Science 6:359–364.

Toda E, Koiso N, Takebayashi A, Ichikawa M, Kiba T, Osakabe K, Osakabe Y, Sakakibara H, Kato N, Okamoto T (2019) An efficient DNA- and selectable-marker-free genome-editing system using zygotes in rice. Nature Plants 5:363–368.

Van Leene J, Boruc J, Jaeger JD, Russinova E, Veylder LD (2011) A kaleidoscopic view of the Arabidopsis core cell cycle interactome. Trends Plant Sci 16:141–50.

Verkest A, Weinl C, Inzé D, De Veylder L, Schnittger A (2005) Switching the Cell Cycle. Kip-Related Proteins in Plant Cell Cycle Control. Plant Physiol 139:1099–1106.

Wang G, Kong H, Sun Y, Zhang X, Zhang W, Altman N, DePamphilis CW, Ma H (2004) Genome-wide analysis of the cyclin family in Arabidopsis and comparative phylogenetic analysis of plant cyclin-like proteins. Plant Physiol 135:1084–1099.

Wang H, Fowke LC, Crosby WL (1997) A plant cyclin-dependent kinase inhibitor gene. Nature 386: 451–452.

Wang K, Liu Y, Li S (2013) Bimolecular Fluorescence Complementation (BIFC) Protocol for Rice Protoplast Transformation. Bio-protocol 3:1–5.

Williams GH, Stoeber K (2012) The cell cycle and cancer. J Pathol 226:352–364.

Xu H, Tsao T (1997) Detection and immunolocalization of glycoproteins of the plasma membrane of maize sperm cells. Protoplasma 198:125–129

Xu H, Zhang J, Zeng J, Jiang L, Liu E, Peng C, He Z, Peng X (2009) Inducible antisense suppression of glycolate oxidase reveals its strong regulation over photosynthesis in rice. J Exp Bot 60:1799–1809.

Yang R, Tang Q, Wang H, Zhang X, Pan G, Wang H, Tu J (2011) Analyses of two rice (Oryza sativa) cyclin-dependent kinase inhibitors and effects of transgenic expression of OsiICK6 on plant growth and development. Ann Bot 107:1087–1101.

Yu X, Zhang X, Zhao P, Peng X, Chen H, Bleckmann A, Bazhenova A, Shi C, Dresselhaus T, Sun MX (2021) Fertilized egg cells secrete endopeptidases to avoid polytubey. Nature 592:433–437.

Zhang S, Tian Z, Li H, Guo Y, Zhang Y, Roberts JA, Zhang X, Miao Y (2019) Genome-wide analysis and characterization of F-box gene family in Gossypium hirsutum L. BMC Genomic 20:1–16.

Zhang X, Zhang X, Gonzalez-Carranza ZH, Zhang S, Miao Y, Liu CJ, Robert JA (2019) F-box proteins in plants. Annual Plant Reviews 2:1–21.

Zheng N, Schulman BA, Song L, Miller JJ, Jeffrey PD, Wang P, Chu C, Koepps DM, Elledge SJ, Pagano M, Conaway RC, Conaway JW, Harper JW, Pavletich NP (2002) Structure of the Cul1-Rbx1-Skp1-F boxSkp2 SCF ubiquitin ligase complex. Nature 416:703–709.

