## Supplemental Information for "A CDKB/KRP/FB3 cell cycle core complex functions in rice gametes and zygotes"

**Supplementary Information**

**Additional file 1**

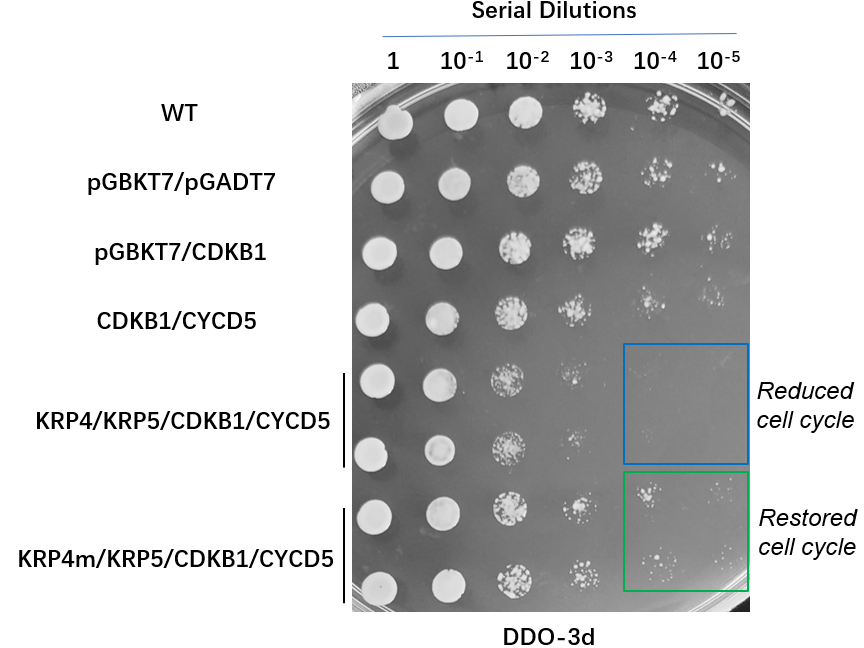

**Fig. S1** Growth of serial dilutions of yeast cells transfected with rice cell cycle core factor genes indicating the reduced cell cycle of yeast cells containing KRP4/KRP5/CDKB1/CYCD5 is partially restored in the cells containing KRP4m/KRP5/CDKB1/CYCD5 (KRP4m: C-terminal truncation).

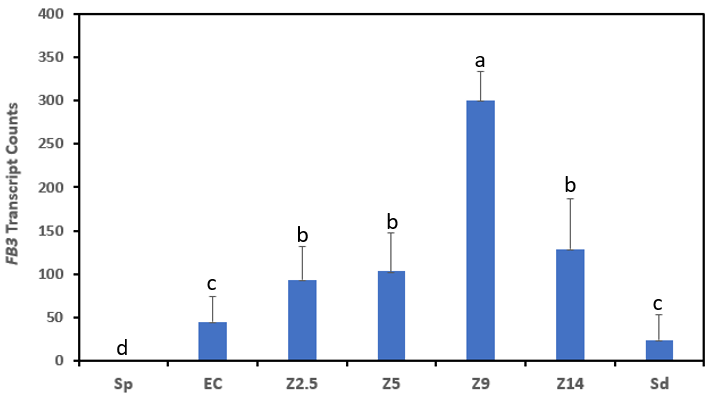

**Fig. S2** RNAseq result of rice F-box gene, *FB3* indicating the preferential expression of *FB3* in rice egg cells and zygotes **(**Anderson et al., 2013; Anderson et al., 2017). Sp, sperm cell; EC, egg cell; Z2.5, zygote at 2.5 hours after pollination (HAP); Z5, zygote of 5 HAP; Z9, zygote of 9 HAP; Sd, seedling. N = 3 biological replicates; values are means and error bars denote the standard deviation. Letters indicate significantly different means determined by one-way ANOVA (with Tukey’s post hoc test;

*P* < 0.001 for a, *P* < 0.01 for b, and *P* < 0.05 for c, compared to Sp).

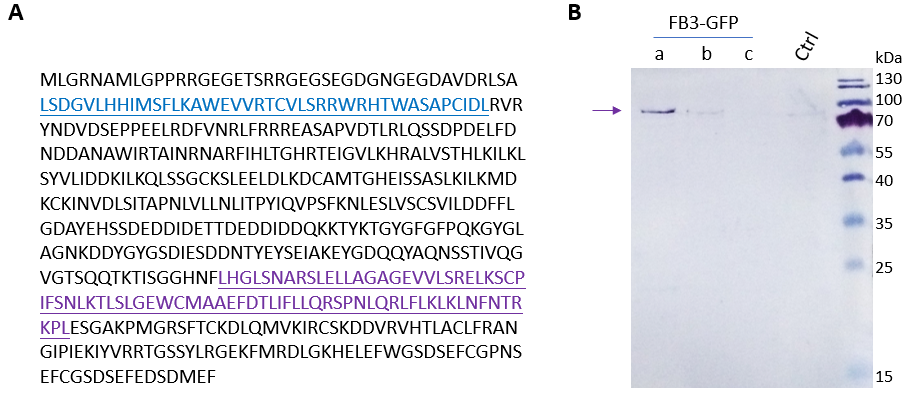

**Fig. S3** FB3 Protein Sequence and Expression in Rice Leaf Protoplasts. **A** Amino acid sequence of FB3 showing the F-box (in blue) and Leucine Rich Region (in purple). **B** Western blot assays of FB3-GFP expressed in rice protoplasts in suspension culture. FB3-GFP (the right, ~89 kDa, arrow pointed) was detected with Anti-GFP mAb from protein extract of rice leaf protoplasts transfected with *FB3-GFP* driven by Maize Ubiqitin promoter. Transfected cells (a, b and c) were cultured in suspension with Chu’s medium plus supplementals for 9 days. The inducer for suspension culture is ß-estrogen (20 µM).

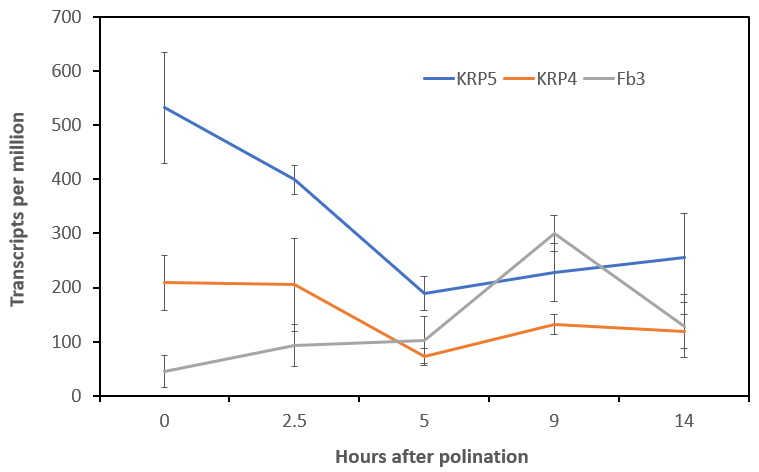

**Fig. S4** Transcription profile of *KRP5, KRP4* and *FB3* in rice egg cells and zygotes with development, based on the RNAseq data **(**Anderson et al., 2013; Anderson et al., 2017)**.** Values are the mean and error bars are the standard deviation from 3 biological replicates.

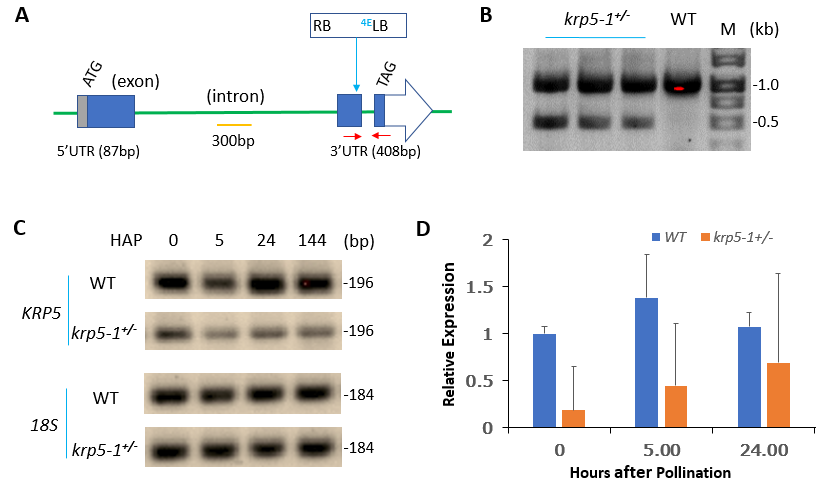

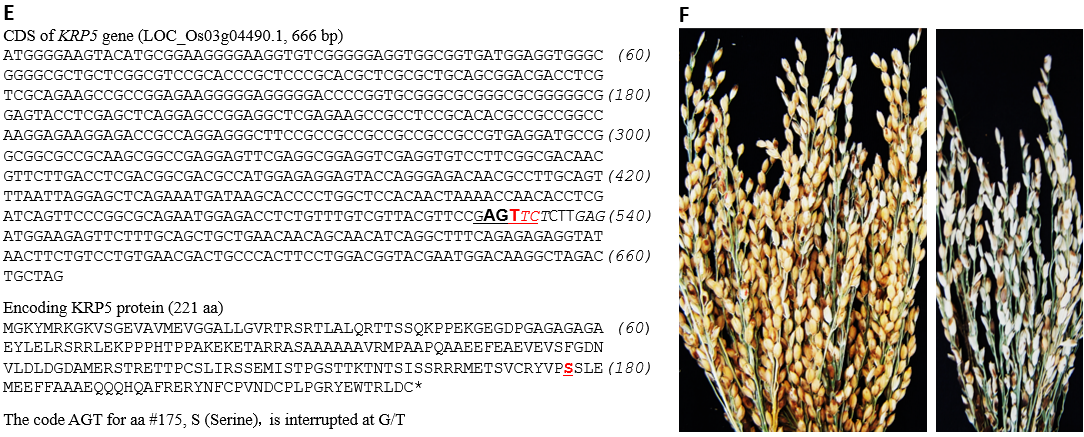

**Fig. S5** Identification and analysis of rice mutant *krp5-1^+/-^* (heterozygous). **A** *KRP5* gene structure (as [*http://rice.plantbiology.msu.edu/ analyses_search.shtml*](http://rice.plantbiology.msu.edu/%20analyses_search.shtml)) and Transfer-DNA (T-DNA) insertion site in the mutant. Red arrows indicate the location of primers for RT-PCR and RT-qPCR; RB and LB, the right and left border of T-DNA; 4E, 4 copies of enhancer elements of CaMV 35S promoter. **B** Agarose gel image of *krp5-1****^+/-^*** genotyping results with primers (Supplementary Table S6) in multiplex PCR: the 1 kb band presents the PCR product from genomic DNA without the T-DNA, and the 0.5 kb band the chimeric product from the T-DNA insertion region; heterozygous individuals are identified by the presence of both products. **C** Gel image of RT-PCR showing dynamics of *KRP5* transcript amounts in wildtype (WT) and mutant ovaries at different developmental stages (HAP, hours after pollination). *18S rRNA* was used as the loading control. **D** Result of RT-qPCR demonstrating reduced *KRP5* transcript abundance in *krp5-1****^+/-^*** ovaries relative to WT at different stages. Shown are means of 3 technical repeats. Error bars are standard deviation. The expression was normalized to WT at 0 HAP. *18S rRNA* gene was used as the internal control. **E** T-DNA insertion site (top panel, shown in red type) within the coding sequence (CDS) of *KRP5* as determined by our DNA sequencing. Predicted amino acid sequence of KRP5 (bottom panel); the red type of color indicates the amino acid after which the T-DNA insertion interrupts the coding sequence. **F** Panicles of WT (the left) and heterozygous mutant (the right) in the T_1_ generation.

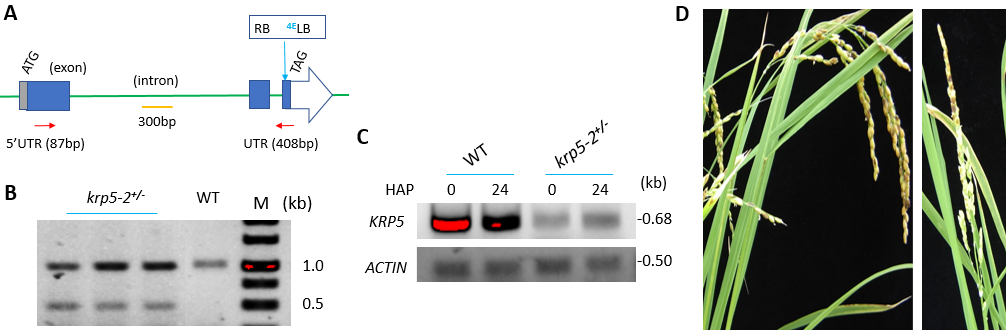

**Fig. S6** Identification and analysis of rice mutant *krp5-2^+/-^* (heterozygous). **A** *KRP5* gene structure and T-DNA insertion site in the mutant confirmed by DNA sequencing; red arrows indicate the location of primers for RT-PCR (C); RB and LB, the right and left border of T-DNA; 4E, 4 copies of 35S promoter enhancer elements. **B** Gel image of *krp5-2****^+/-^*** genotyping with specific primers (Supplementary Table S6) in multiplex PCR: the 1 kb band presents the PCR product from genomic DNA, and the 0.5 kb band the chimeric product from T-DNA insertion region; the heterozygous individual plants are identified by the presence of both products. **C** Gel image of RT-PCR showing reduced KRP5 transcription level in *krp5-2****^+/-^*** ovaries at 0 and 24 HAP, and *ACTIN* used as the internal control. **D** Representative panicle of WT (left) and *krp5-2****^+/-^*** (right) in T_1_ generation.

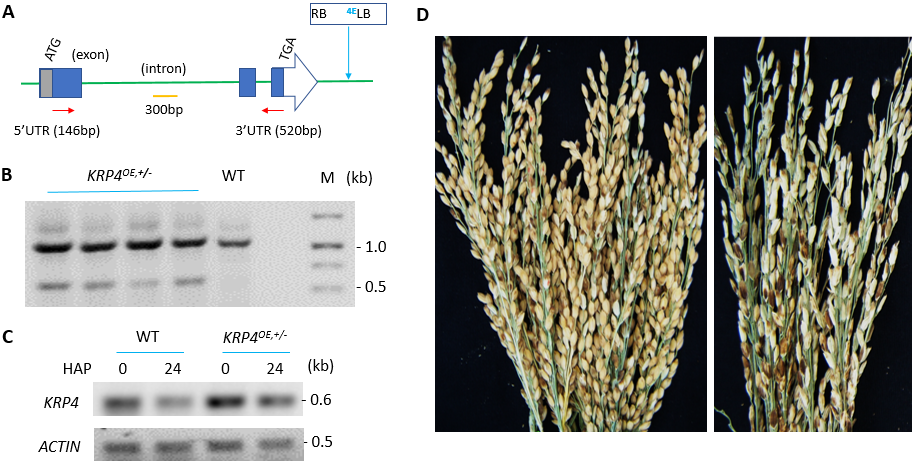

**Fig. S7** Identification and analysis of rice mutant *KRP4-D1^OE/+^* (heterozygous). **A** *KRP4* gene structure (as [*http://rice.plantbiology.msu.edu/analyses_search.shtml*](http://rice.plantbiology.msu.edu/analyses_search.shtml)), T-DNA insertion site determined by DNA sequencing. Red arrows indicate the location of primers for RT-PCR (c); 4E, 4 copies of *35S* promoter enhancer elements; *KRP4-D1^OE/+^*, the mutant with overexpression of *KRP4*. **B** Agarose gel image of *KRP4-D1^OE/+^* genotyping results with primers (SupplementaL Table S6) in multiplex PCR: the 1 kb band presents the PCR product from genomic DNA without T-DNA, and the 0.5 kb band the chimeric product from the T-DNA insertion region; the heterozygous individuals are identified by the presence of both products. **C** Gel image of RT-PCR demonstrating enhanced expression of *KRP4* gene in *KRP4-D1^OE/+^* ovaries at and 24 HAP, and *ACTIN* was used as the internal control. **D** Panicles of WT (the left) and *KRP4-D1^OE/+^* (the right) in T_1_ generation.

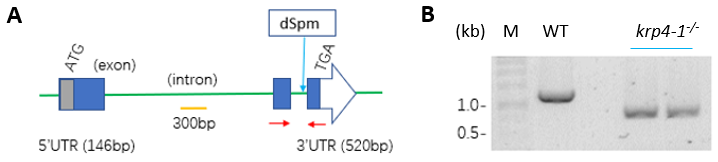

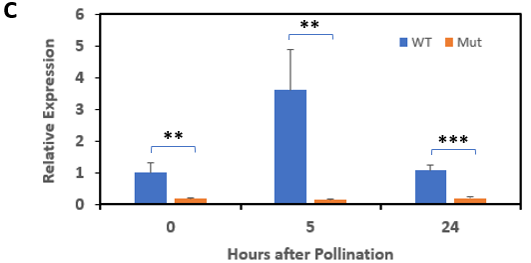

**Fig. S8** Identification and analysis of rice mutant *krp4-1^-/-^* (homozygous). **A** *KRP4* gene structure and the location (red arrows) of primers for RT-qPCR (C) and the insertion site of transposon dSpm within intron upstream of the STOP, confirmed by DNA sequencing. **B** Agarose gel image of *krp4-1****^-/-^*** genotyping with primers (Supplementary Table S6) in multiplex PCR: the 1.1 kb band presents the PCR product from genomic DNA without T-DNA, and the 0.7 kb band shows the chimeric product from the T-DNA insertion region; the homozygous individuals are identified by the absence of 1.1 kb product. **C** RT-qPCR demonstrating reduced expression of *KRP4* gene in *krp4-1* ***^-/-^*** ovaries at 0, 5 and 24 HAP. *18S rRNA* gene used as the internal control. The expression was normalized to WT at 0 HAP. Values are means of 3 biological replicates. Error bars are standard deviation. A t-test was performed to indicate significant differences at *P* < 0.001 (***) and *P* < 0.01 (**).

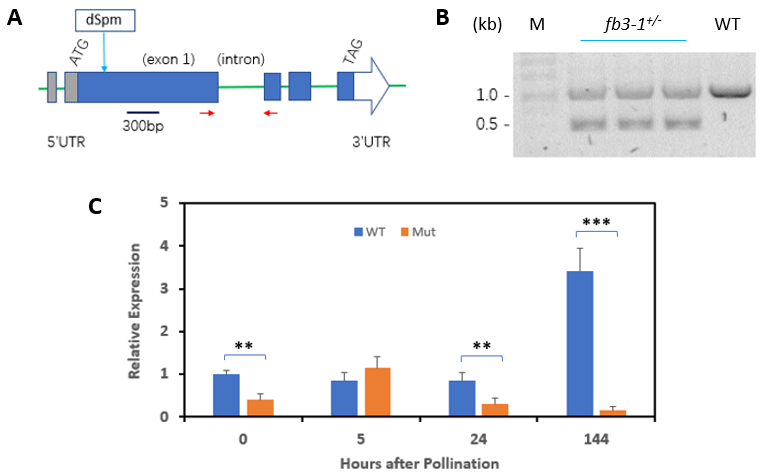

**Fig. S9** Identification and analysis of rice mutant *fb3-1^+/-^* (heterozygous). **A** *FB3* gene structure with the location (red arrows) of primers for RT-qPCR and the insertion site of transposon dSpm within the Exon 1, determined by DNA sequencing. **B** Agarose gel image of *fb3-1****^+/-^*** genotyping with primers (Supplementary Table S6) in multiplex PCR: the 1.0 kb band presents the PCR product from genomic DNA without T-DNA, and the 0.5 kb band shows the chimeric product from the T-DNA insertion region; the heterozygous individuals are identified by the presence of both products. **C** RT-qPCR demonstrating dynamics of *FB3* transcript amount in ovaries at 0, 5, 24 and 144 HAP and *18S rRNA* was used as the internal control. The relative expression was normalized with WT at 0 HAP. Values are means of 3 biological replicates. Error bars are standard deviation. The t-test was performed to indicate significant differences at *P* < 0.001 (***) and *P* < 0.01 (**).

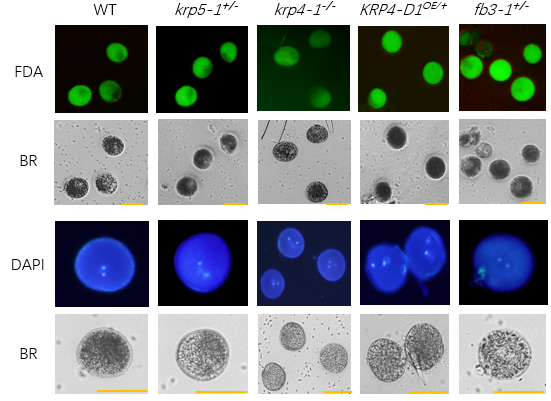

**Fig. S10** The pollen of rice mutants stained with FDA and DAPI. Staining with FDA is to evaluate the pollen viability and staining with DAPI to observe the morphology of sperm cells within pollen grains. BR, pollen images in bright field. Scale bars: 50 µm. N = ~ 200 pollen grains per line; no significant differences were observed in the viability and two sperm cells were present in each grain.

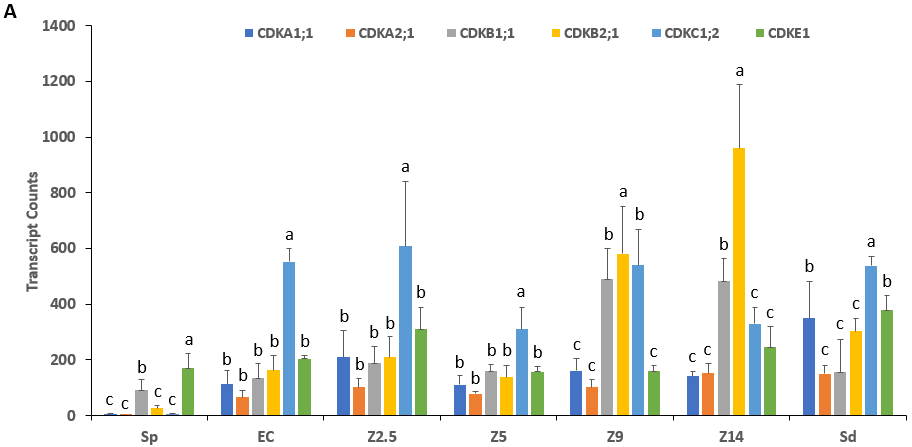

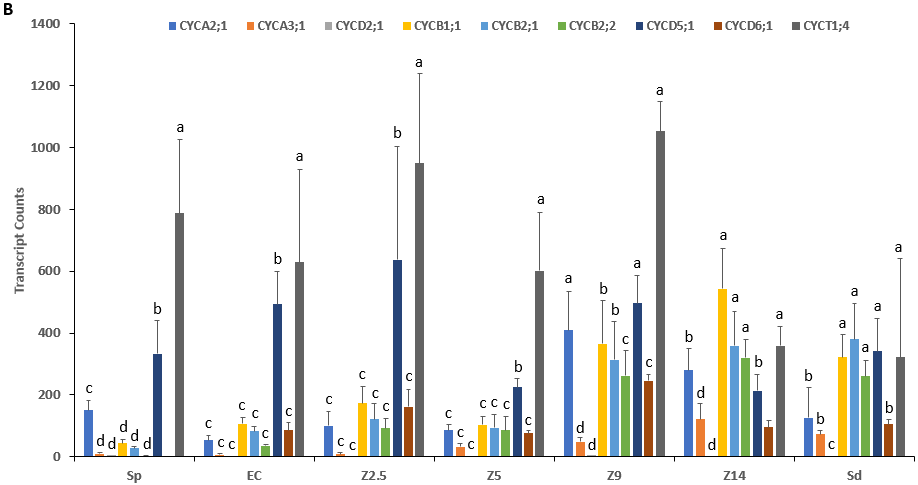

**Fig. S11** RNAseq results of 6 rice *CDK* genes and 9 rice *CYC* genes (locus numbers are listed in Table S1; based on Anderson et al., 2013 and Anderson et al., 2017). **A** *CDKB1;1, CDKB2;1* and *CDKC1;2* are preferentially expressed in rice egg cells and zygotes. **B** *CYCD5;1* and *CYCT1;4* are preferentially expressed in rice egg cells and zygotes. Sp, sperm cell; EC, egg cell; Z2.5, zygote at 2.5 hours after pollination (HAP); Z5, zygote of 5 HAP; Z9, zygote of 9 HAP; Z14, zygote of 14 HAP; Sd, seedling. N = 3 biological replicates; columns indicate mean values and error bars denote standard deviation. Different letters indicate significantly different groups determined by one-way ANOWA (*P* < 0.05).

**Additional file 2**

**Table S1** Rice cell cycle gene length, expression in rice flowers, and cloning success

| Name | Locus **(*LOC_Os*)** | CDS ***(bp)*** | RT-PCR^39^ | RNA-seq^1^ | RNAseq^2^ | RT-PCR^3^ | Cloned |
| --- | --- | --- | --- | --- | --- | --- | --- |
| *KRP1* | *02g52480* | 798 |  | 13 - 27 | 24.3 ± 3.8 | +++ | Y |
| *KRP2* | *06g11050* | 1233 |  | 0 | 0.3 ± 0.5 |  |  |
| *KRP3* | *01g40030* | 678 |  | 0 - 2 | 0 |  |  |
| *KRP4* | *10g33310* | 585 | + | 3 - 65 | 71.7 ± 13.0 | ++++ | Y |
| *KRP5* | *03g04490* | 666 | + | 1 - 32 | 189.3 ± 25.2 | ++++ | Y |
| *KRP6* | *01g37740* | 615 |  | 0 - 1 | 0 |  |  |
| *CDKA1;1* | *03g02680* | 999 | +++ | 7 - 46 | 110.7 ± 27.6 | + | Y |
| *CDKA2;1* | *02g03060* | 1131 | +++ | 2 - 14 | 78.3 + 7.7 | ++ | Y |
| *CDKB1;1* | *01g67160* | 912 | ++ | 3 - 26 | 160.7 ± 17.9 | ++++ | Y |
| *CDKB2;1* | *08g40170* | 981 | +++ | 5 - 55 | 137.7 ± 33.9 | +++ | Y |
| *CDKC1;1* | *05g32360* | 1560 | + | 17 - 18 | 40.6 ± 10.5 |  |  |
| *CDKC1;2* | *01g72790* | 1542 | +++ | 38 - 58 | 310.7 ± 64.5 | +++ | Y |
| *CDKC3;1* | *08g35220* | 975 | ++ | 6 - 12 | 4.3 ± 1.1 |  |  |
| *CDKD1* | *05g32600* | 1275 | ++ | 13 - 17 | 24.7 ± 5.3 |  |  |
| *CDKE1* | *10g42950* | 1428 | +++ | 15 - 35 | 158.3 ± 13.9 | +++ | Y |
| *CYCA2;1* | *12g31810* | 1473 | +++ | 6 - 8 | 13.1 ± 2.4 | ++++ | Y |
| *CYCA3;1* | *03g41100* | 1122 | + | 5 - 13 | 8.4 ± 1.3 | ++ | Y |
| *CYCA3;2* | *12g39210* | 1158 |  | 0 - 8 | 9.2 ± 2.7 | + | Y |
| *CYCB1;1* | *01g59120* | 1350 | +++ | 24 - 33 | 17.1 ± 5.2 | ++++ | Y |
| *CYCB1;3* | *05g41390* | 1350 |  | 12 - 20 | 12.2 ± 5.0 |  |  |
| *CYCB2;1* | *04g47580* | 1263 | +++ | 12 - 16 | 16.9 ± 7.9 | +++ | Y |
| *CYCB2;2* | *06g51110* | 1260 | +++ | 8 - 16 | 15.1 ± 7.6 | +++ | Y |
| *CYCD1;1* | *08g32540* | 1065 |  | 8 - 10 | 0.8 ± 0.4 |  |  |
| *CYCD1;2* | *06g12980* | 1092 |  | 0 | 0 |  |  |
| *CYCD2;1* | *09g21450* | 690 | +++ | 6 - 15 | 3.7 ± 1.2 | + | Y |
| *CYCD3;1* | *09g02360* | 1095 |  | 10 - 23 | 10.6 ± 3.3 | ++ | Y |
| *CYCD4;1* | *07g42860* | 1071 | +++ | 8 - 14 | 24.4 ± 3.9 |  |  |
| *CYCD4;2* | *06g11410* | 1029 | ++ | 8 - 21 | 13.2 ± 5.0 |  |  |
| *CYCD4;3* | *08g37390* | 1152 |  | 8 - 18 | 12.5 ± 2.7 |  |  |
| *CYCD4;4* | *09g29100* | 1071 |  | 10 - 15 | 10.5 ± 4.2 |  |  |
| *CYCD4;5* | *03g27420* | 1218 |  | 0 | 0.4 ± 0.1 |  |  |
| *CYCD5;1* | *03g10650* | 1038 | +++ | 6 - 61 | 224 ± 22.7 | +++ | Y |
| *CYCD5;2* | *03g42070* | 1104 |  | 1 - 8 | 12.7 ± 5.3 |  |  |
| *CYCD5;3* | *12g39830* | 1098 |  | 0 - 19 | 6.2 ± 2.3 |  |  |
| *CYCD6;1* | *07g37010* | 963 | ++++ | 43 - 48 | 14.2 ± 1.3 | ++ | Y |
| *CYCD7;1* | *11g47950* | 963 | ++ | 0 | 0 |  |  |
| *CYCH1;1* | *03g52750* | 993 | ++ | 6 - 7 | 18.2 ± 5.9 |  |  |
| *CYCT1;1* | *02g04010* | 1341 | +++ | 25 - 30 | 16.4 ± 1.5 |  |  |
| *CYCT1;4* | *12g30020* | 1632 | +++ | 22 - 34 | 601 ± 153.8 | ++++ | Y |

1. RNA-seq of rice anther and pistil in FPKM, MSU (<http://rice.plantbiology.msu.edu>)

2. RNA-seq of rice zygotes at 5 HAP (hours after pollination) in TPM (transcripts per million) **(**Anderson et al.,

2013; Anderson et al., 2017).

3. RT-PCR conducted in this work. The number of +’s indicates the signal intensity of RT-PCR product in DNA gel.

4. Y indicates that the gene was cloned.

**Table S2** Locus numbers of F-box genes expressed in rice zygotes

| **Name** | **Locus #^1^** | **CDS (bp)** | **RNAseq (TMP)^2^** | **RT-PCR^3^** | **Cloned^3^** |
| --- | --- | --- | --- | --- | --- |
| *FB1* | *LOC_Os03g43990* | 1875 | 235.7 ± 41.3 | ++ | Y |
| *FB2* | *LOC_Os12g40860* | 1722 | 210.3 ± 7.0 | ++ | Y |
| *FB3* | *LOC_Os08g09750* | 1656 | 103.3 ± 35.8 | + | Y |
| *FB4* | *LOC_Os06g40360* | 1995 | 185.7 ± 10.1 |  |  |
| *FB5* | *LOC_Os05g37690* | 1794 | 230.3 ± 52.2 |  |  |
| *FB6* | *LOC_Os05g35110* | 1131 | 108 ± 30.3 | ++ | Y |
| *FB7* | *LOC_Os04g32460* | 1728 | 496.7 ± 42.7 |  |  |

1. Rice genomic database, MSU: http://rice.plantbiology.msu.edu.

2. RNA-seq in rice zygotes at 5 HAP (hours after pollination) in TPM (transcripts per million) **(**Anderson et al.,

2013; Anderson et al., 2017).

3. RT-PCR and cloning done in this work: +, signal intensity of RT-PCR product in DNA gel; Y, successfully

cloned.

**Table S3** Five rice mutant lines*

| **Mutant** | **Locus (*LOC*_)** | **Line #** | **Line name** | **WT** | **Vectors** | **Reference** |
| --- | --- | --- | --- | --- | --- | --- |
| *krp5-1****^+/-^*** | *Os03g04490* | 05609 | PFG_3A-05609.R | DJ | pGA2715 | Jeong et al., 2006 |
| *krp5-2****^+/-^*** | *Os03g04490* | 18132 | PFG_1A-18132.L | HY | pGA2715 | Jeong et al., 2002 |
| *KRP4-D1^OE^*^/^**^+^** | *Os10g33310* | 06157 | PFG_3A-06157.L | DJ | pGA2715 | Jeong et al., 2006 |
| *krp4-1****^-/-^*** | *Os10g33310* | 6385A | RdSpm6385A_3.1 | Nip | pdSpm-imm.Spm | Kumar et al., 2005 |
| *fb3-1***^+/-^** | *Os08g09750* | 2792 | RdSpm2792_3.1 | Nip | pdSpm-imm.Spm | Kumar et al., 2005 |

*** Selected from Rice Functional Genomic Express Database *(*[*http://signal.salk.edu/cgi-bin/RiceGE*](http://signal.salk.edu/cgi-bin/RiceGE)*).*

DJ, Dongjin; HY, Huayoung, Nip, Nipponbare

**Table S4** Analysis of seed-yield related traits in rice mutant lines (T_2_)

and wild type (values are mean ± standard deviation).

| **WT/Mutants** | **Panicles/plant** | **Seeds/panicle** | **Seed-set rate (%)** | **1000-seedweight (g)** |
| --- | --- | --- | --- | --- |
| WT (DJ) | 7.7 ± 0.7 | 54.3 ± 10.5 | 89 ± 7.5 | 24.2 ± 0.4 |
| ***krp5-1^+/-^*** | 7.3 ± 2.1 | 47.3 ± 11.2 | 33.1 ± 9.4^**^ | 21.3 ± 0.2^**^ |
| ***KRP4-D1^OE/^*^+^** | 9.3 ± 0.6 | 38.3 ± 8.7 | 34 ± 8.7^**^ | 20.8 ± 0.5^**^ |
| WT (HY) | 7.7 ± 1.5 | 42 ± 6 | 91.7 ± 2.3 | 24.2 ± 0.4 |
| ***krp5-2^+/-^*** | 5.3 ± 1.5 | 48.6 ± 9 | 28 ± 6.9^**^ | 19.2 ± 0.8^**^ |
| WT (Nip) | 7.3 ± 1.5 | 63.3 ± 5 | 86.6 ± 5.5 | 22.3 ± 0.2 |
| ***krp4-1^-/-^*** | 8 ± 1 | 54.3 ± 4.7 | 56.1 ± 15.8^**^ | 22 ± 0.1^*^ |
| ***fb3-1*^+/-^** | 8.7 ± 1.5 | 69.7 ± 14.6 | 46.1 ± 9.9^**^ | 21.9 ± 0.7^**^ |

*, Significant difference *versus* WT in t-test at *P* < 0.05.

**, Significant difference *versus* WT in t-test at *P* < 0.01.

WT: DJ, Dongjin; HY, Huayoung; Nip, Nipponbare

**Table S5** Primers used in this study

| **Primers** | **Sequences (5’-3’)** | **Purposes** |
| --- | --- | --- |
| KRP1-RT-F | ATGGGCAAGTACATGAGGAAG | For |
| KRP1-RT-R | TCAGCTTCGGCTGCTGAC C | RT-PCR |
| KRP2-RT-F | GACTGAATTCATGGGGAAGAAGAAGAAGC |  |
| KRP2-RT-R | CTGAGGATCCTCAGTGAACGCGGAGGCG |  |
| KRP3-RT-F | GTCACATATGATGGGCAAGTACCTCAGGAG |  |
| KRP3-RT-R | CAGTCTCGAGTCACATGTCAGTGACATCCTT |  |
| KRP4-RT-F | GAGGCCGAATTCATGGGCAAGTACATGCGCAA |  |
| KRP4-RT-R | GTCGACGGATCCTCAGTCTAGCTTGACCCATTC |  |
| KRP5-RT-F | GAGGCCGAATTCATGGGGAAGTACATGCGGAAG |  |
| KRP5-RT-R | GTCGACGGATCCCTAGCAGTCTAGCCTTGTCCA |  |
| KRP6-RT-F | GTGCCCCGGGATGCTCGGGAGGAACGCCA |  |
| KRP6-RT-R | CGTGCCTCGAGCTCCTCCGGCGGCTCGCT |  |
| FB1-RT-F | CGTGCATGGCGTCGCTGCCCAAGC |  |
| FB1-RT-R | CGTGCTCAATCAAGATGAACGGTGC |  |
| FB2-RT-F | CGTGCATGGCCTCCTCCTCCGACG |  |
| FB2-RT-R | CGTGCTTAGCTCAGGTGAACGGTGC |  |
| FB3-RT-F | CGTGCCATGCTCGGGAGGAACGCCA |  |
| FB3-RT-R | CGTGCTCAGAACTCCATATCTGAATC |  |
| FB4-RT-F | CTGCATGCCTCCGTTCCACGATT |  |
| FB4-RT-R | GTCTCTAAGCAAGGATGTCGCA |  |
| FB5-RT-F | CTGCATGGGAGGGGAGGCACCG |  |
| FB5-RT-R | GTCTTCACGCAGGATACAAAGGA |  |
| FB6-RT-F | CTGCATGGTTAGTGGGAGGTC |  |
| Fb6-RT-R | GTCTTCAGTATGCGTGGCTAGG |  |
| FB7-RT-F | CTGCATGACGTACTTCCCGGAG |  |
| FB7-RT-R | GTCTCTATAGGATTTTAACAAAATTT |  |
| OsActin-RT-F | CTTTGCCTTGAGATGGACGC |  |
| OsActin-RT-R | TCACCAGGACACCACCAAAC |  |
| 18S-RT-F | TTAGGCCACGGAAGTTTGAG |  |
| 18S-RT-R | GCATTCCTCGTTGAAGACCA |  |
| KRP4-qRT-F | CCGATACGATTAGCACCCCT | For |
| KRP4-qRT-R | AGGGCAGTCATTCACAGGAT | RT-qPCR |
| KRP5-qRT-F | ACCAACACCTCGATCAGTTCC |  |
| KRP5-qRT-R | CTAGCAGTCTAGCCTTGTCC |  |
| Fb3-qRT-F | TGGAGATCAGCAGTATGCCC |  |
| Fb3-qRT-R | GCAGCTTTTCAACTCCCTAGAA |  |
| 18S-qRT-F | TTAGGCCACGGAAGTTTGAG |  |
| 18S-qRT-R | GCATTCCTCGTTGAAGACCA |  |
| KRP1-Y2Hbait-F-EcoRI | GACTGAATTCATGGGCAAGTACATGAGGAAG | For |
| KRP1-Y2Hbait-R-BamHI | CTGAGGATCCTCAGCTTCGGCTGCTGACC | Y2H assay |
| KRP4-F-EcoRI | GAGGCCGAATTCATGGGCAAGTACATGCGCAA |  |
| KRP4-R-BamHI | GTCGACGGATCCTCAGTCTAGCTTGACCCATTC |  |
| KRP5-F-EcoRI | GAGGCCGAATTCATGGGGAAGTACATGCGGAAG |  |
| KRP5-R-BamHI | GTCGACGGATCCCTAGCAGTCTAGCCTTGTCCA |  |
| FB1-F-NdeI | CGTGCCATATGATGGCGTCGCTGCCCAAGC | For |
| FB1-R-XhoI | CGTGCCTCGAGTCAATCAAGATGAACGGTGC | Y2H assay |
| FB2-F-NdeI | CGTGCCATATGATGGCCTCCTCCTCCGACG |  |
| FB2-R-XhoI | CGTGCCTCGAGTTAGCTCAGGTGAACGGTGC |  |
| FB3-F-NdeI | CGTGCCATATGATGCTCGGGAGGAACGCCA |  |
| FB3-R-XhoI | CGTGCCTCGAGTCAGAACTCCATATCTGAATC |  |
| FB6-F-EcoRI | CTGCGAATTCATGGTTAGTGGGAGGTC |  |
| FB6-R-BamHI | GTCTGGATCCTCAGTATGCGTGGCTAGG |  |
| CDKA1;1-Y2Hprey-F-EcoRI | CTGCGAATTCATGGAGCAGGTGAGCGCCT |  |
| CDKA1;1-Y2Hprey-R-BamHI | GTCTGGATCCTCATTGTACCATCTCAAGGT |  |
| CDKA2;1-Y2Hprey-F-EcoRI | CTGAGAATTCATGCCACAAGCCCAACCCA |  |
| CDKA2;1-Y2Hprey-R-BamHI | GACTGGATCCCTACGCCACTTCCAGGTCC |  |
| CDKB1;1-Y2Hprey-F-NdeI | GTCACATATGATGGAGAAGTACGAGAAGCTG |  |
| CDKB1;1-Y2Hprey-R-XhoI | CAGTCTCGAGCTAGAACGTGGACTTGTCGA |  |
| CDKB2;1-Y2Hprey-F-EcoRI | CTGAGAATTCATGGCGGCGCTCCACCAC |  |
| CDKB2;1-Y2Hprey-R-BamHI | GACTGGATCCTCAGTAGAGCTCCTTGTTCAC |  |
| CDKC1;2-Y2Hprey-F-NdeI | GTCACATATGATGGCGGTGGCGGCGCC |  |
| CDKC1;2-Y2Hprey-R-EcoRI | CTGAGAATTCCTATTGCCAGTTTCCATACTG |  |
| CDKE1-Y2Hprey-F-NdeI | GTCACATATGATGGGGGACGGCCGCGT |  |
| CDKE1-Y2Hprey-R-EcoRI | CTGAGAATTCTCAGAACCGCCTGTTCTGTT |  |
| CYCA2;1-Y2Hprey-F-NdeI | CGTGCCATATGATGGCTGGAAGGAAGGAAAAT |  |
| CYCA2;1-Y2Hprey-R-XhoI | CGTGCCTCGAGTACGCTGAAGAGTGACTGCC |  |
| CYCA3;1-Y2Hprey-F-NdeI | CGTGCCATATGATGGCCGGCAAGGAGAACG |  |
| CYCA3;1-Y2Hprey-R-XhoI | CGTGCCTCGAGCTACTCGTTTAGGTCTTCGAA |  |
| CYCA3;2-Y2Hprey-F-EcoRI | CTGCGAATTCATGGCTGACAAGGAGAACTC |  |
| CYCA3;2-Y2Hprey-R-BamHI | GTCTGGATCCCTACTCTGTTAAATCTTGGAGG |  |
| CYCB1;1-Y2Hprey-F-EcoRI | CTGCGAATTCATGGCGACACGCAGCCAG |  |
| CYCB1;1-Y2Hprey-R-BamHI | GTCTGGATCCTTATTTGCAGAGTTCCACTGC |  |
| CYCB2;1-Y2Hprey-F-NdeI | CGTGCCATATGATGGATAGGGCGAGCGAGA |  |
| CYCB2;1-Y2Hprey-R-XhoI | CGTGCCTCGAGTCAACAAGGCTGCTTCTGAAG |  |
| CYCB2;2-Y2Hprey-F-NdeI | CGTGCCATATGATGGAGAACATGAGATCTGAG |  |
| CYCB2;2-Y2Hprey-R-XhoI | CGTGCCTCGAGTTACAGTGCCACGCTCTTGA |  |
| CYCD2;1-Y2Hprey-F-NdeI | CGTGCCATATGATGCCGGCCGACGACGAC |  |
| CYCD2;1-Y2Hprey-R-XhoI | CGTGCCTCGAGTCACCCAGCAAAAACATCTGT |  |
| CYCD5;1-Y2Hprey-F-EcoRI | CTGAGAATTCATGGGGGACGCCTCGGC |  |
| CYCD5;1-Y2Hprey-R-BamHI | GACTGGATCCCTACTGGCGCTGAGGCGA |  |
| CYCD6;1-Y2Hprey-F-EcoRI | CTGAGAATTCATGGACATGGCGACGGGG |  |
| CYCD6;1-Y2Hprey-R-BamHI | GACTGGATCCTCACTCATCAGGGCCGCC |  |
| CYCT1;4-Y2Hprey-F-NdeI | GTCACATATGATGGCTATGATGCCAAGTGAT |  |
| CYCT1;4-Y2Hprey-R-XhoI | CAGTCTCGAGCTAACCTTCAGGCCGAGGC |  |
| KRP4-F-KpnI | TGTGGTACCATGGGCAAGTACATGCGCAA | For BiFC |
| KRP4-R-SmaI | TGTCCCGGGCCGTCTAGCTTGACCCATTCAAA | and |
| KRP5-F-EcoRI | GTCGAATTCTATGGGGAAGTACATGCGGAAG | cellular |
| KRP5-R-SalI | TGCGTCGACCTAGCAGTCTAGCCTTGTCCA | localization |
| UBIQprom-F-AgeI | TGTACCGGTCTGCAGTGCAGCGTGACCC | assays |
| UBIQprom-R-HindIII | TGCAAGCTTCTGCAGAAGTAACACCAAACAA |  |
| KRP5-F-EcoRI | GTCGAATTCTATGGGGAAGTACATGCGGAAG |  |
| KRP5-R-SalI | TGCGTCGACGCAGTCTAGCCTTGTCCATTC | For BiFC |
| KRP5prom-F-AgeI | CGTGTACCGGTGGTGACACATCGGTGGAGC | and |
| KRP5prom-R-SmaI | CGTGTCCCGGGGGCCAAGGCGCGCGCGG | cellular |
| CDKB1-F-PstI | GTCACTGCAGATGGAGAAGTACGAGAAGCTG | localization |
| CDKB1-R-KpnI | CAGTGGTACCCTAGAACTGGGACTTGTCGA | assays |
| CDKB1-cEYFP-F | CATCGGGGATTCGGGATCGATGGAGAAGTACGAGAAGC  TG |  |
| CDKB1-cEYFP-R | CTATTATGGGATTTATGGGTGCTGCGGCCGCGTCACTGG  A |  |
| CDKB1prom-F | CTAAAACCAAAATCCAGTGACGCGTGGTTTTGTGAAAC  TACACTAC |  |
| CDKB1prom-R | CAGCTTCTCGTACTTCTCCATCGATCCCGAATCCCCGA  TG |  |
| UBIQprom-KRP4-cEYFP-F | CATAATAGCTGTTTGCCAACTGCAGTGCAGCGTGACC |  |
| UBIQprom-KRP4-cEYFP-R | CTATTATGGGATTTATGGGTGCTGCGGCCGCGTCACTGG  A |  |
| FB3-F-XhoI | TGTCTCGAGCTATGCTCGGGAGGAACGCCAT |  |
| FB3-R-HindIII | TGCAAGCTTCGGAACTCCATATCTGAATCCTC |  |
| FB3 (5’-300bp)-F-XhoI | TGTCTCGAGCTATGCTCGGGAGGAACGCCAT |  |
| FB3-(5’-300bp)-R-HindIII | TGCAAGCTTCGCTCCTCCGGCGGCTCGCT |  |
| FB3-(3’-1.4kb)-F-XhoI | TGTCTCGAGCTCTGCGCGATTTCGTGAACC |  |
| FB3-(3’-1.4kb)-R-HindIII | TGCAAGCTTCGGAACTCCATATCTGAATCCTC |  |
| FB3prom-F-AgeI-NotI | CGTGTACCGGTGCGGCCGCAGCTCCGATTCATCTATAG  TC |  |
| FB3prom-R-EcoRV | CGTGTGATATCTCCGCCGAACAGGCTGCG |  |
| CDKB1;1-F-XhoI | GTCACTCGAGATGGAGAAGTACGAGAAGCTG | For |
| CDKB1;1-R-EcoRI | CAGTGAATTCGAACTGGGACTTGTCGAGG | mcBiFC |
| KRP4-F-EcoRI | GTCGAATTCTATGGGCAAGTACATGCGCAA | assay |
| KRP4-R-SalI | TGCGTCGACGTCTAGCTTGACCCATTCAAA |  |
| UBIQprom-KRP4-nVenus-F | TAAAACCAAAATCCAGTGACGCCTGCAGTGCAGCGTG  ACC |  |
| UBIQprom-KRP4-nVenus-R | GCTCCACCGATGTGTCACCGCGTCACTGGATTTTGGTTT  TA |  |
| KRP5prom-F-AgeI | CGTGTACCGGTGGTGACACATCGGTGGAGC |  |
| KRP5prom-R-XhoI | CAGTCTCGAGGGCCAAGGCGCGCGCG |  |
| KRP5prom-KRP5-nCerullean-F | TAAAACCAAAATCCAGTGACGCGGTGACACATCGGTG GAGC |  |
| KRP5prom-KRP5-nCerulean-R-NotI | TACGGCGCGCCGCGGCCGCGTCACTGGATTTTGGTTTT  A |  |
| FB3-HA-F-XhoI | TGTCTCGAGCTATGCTCGGGAGGAACGCCAT | For FB3 |
| FB3-HA-R-HindIII | TGAAGCTTCGAGCGTAATCTGGTACGTCGTATGGGTAC TCCATGAACTCCATATCTGAATCCTC | expression  in rice |
| FB3-F-SpeI | CGTGCACTAGTATGCTCGGGAGGAACGCC | protoplasts |
| FB3-R-SpeI | CGTGCACTAGTTCAGAACTCCATATCTGAATC |  |
| FB3-EYFP-F-ApaI | CGTGTGGGCCCCTATGGTGAGCAAGGGCGAG |  |
| FB3-EYFP-R-SpeI | CGTGCACTAGTTCAGAACTCCAT TCTGA TC |  |
| KRP5mut05609-F | AGATGCAACCTCGTGAAAGG | Genotyping |
| KRP5mut05609-R1 | AGCAGGTTTATGTGGCCAAG | rice |
| KRP5mut05609-R2 | AGTTTTCGCGATCCAGACTG | mutants |
| KRP5mut18132-F | TCGTCCAAAGCTGAATGTTG |  |
| KRP5mut18132-R1 | GGTAAACAGAAGGCAGCTGC |  |
| KRP5mut18132-R2 | ACCTCCGGAATCACTGCAA | Genotyping |
| KRP4mut06157-F1 | TTCTTGCAGGAATACCAGGG | rice |
| KRP4mut06157-F2 | CCTGTGCCATTATTGCTACC | mutants |
| KRP4mut06157-R | CCCACTAAGCCTGAGCAGAG |  |
| KRP4mut6385A-F | AAACAAGAAAAATGCACCGG | Genotyping |
| KRP4mut6385A-R1 | ATCTCACCACCCACCTCAAG | rice |
| KRP4mut6385A-R2 | GAGCGTCCATTTTAGAGTGAC | mutants |
| FB3mut2792A-F | AATGAGGCCTCGTTTGAATG | Genotyping |
| FB3mut2792A-R1 | GCGCAGTCCTTAAGATCCAG | rice |
| FB3mut2792A-R2 | GAGCGTCCATTTTAGAGTGAC | mutants |
